# Leafy and Weedy Seadragon Genomes Connect Genic and Repetitive DNA Features to the Extravagant Biology of Syngnathid Fishes

**DOI:** 10.1101/2021.09.24.461757

**Authors:** Clayton M. Small, Hope M. Healey, Mark C. Currey, Emily A. Beck, Julian Catchen, Angela S. P. Lin, William A. Cresko, Susan Bassham

## Abstract

Seadragons are a remarkable lineage of teleost fishes, and they are members of the family Syngnathidae renowned for having evolved male pregnancy. Comprising three known species, seadragons are widely recognized and admired for their fantastical body forms and coloration, and their specific habitat requirements have made them flagship representatives for marine conservation and natural history interests. Until recently, a gap has been the lack of significant genomic resources for seadragons. We have produced gene-annotated, chromosome-scale genome models for the leafy and weedy seadragon to advance investigations into evolutionary innovation and elaboration of morphological traits in seadragons as well as their pipefish and seahorse relatives. We identified several interesting features specific to seadragon genomes, including divergent non-coding regions near a developmental gene important for integumentary outgrowth, a high genome-wide density of repetitive DNA, and recent expansions of transposable elements and a vesicular trafficking gene family. Surprisingly, comparative analyses leveraging the seadragon genomes and additional syngnathid and outgroup genomes revealed striking, syngnathid-specific losses in the family of fibroblast growth factors (FGFs), which likely involve re-organization of highly conserved gene regulatory networks in ways that have not previously been documented in natural populations. The resources presented here serve as important tools for future evolutionary studies of developmental processes in syngnathids and will be a key resource for conservation studies of the extravagant seadragons and their relatives.

## INTRODUCTION

Seadragons are phenotypic outliers in an already exceptional clade of teleost fishes (Family Syngnathidae) that also includes seahorses and pipefishes. For this reason, seadragons are often a colorful, flagship group in discussions of adaptation and evolutionary innovation. Seadragons demonstrate strikingly derived characters compared to their pipefish and seahorse relatives, including “leafy appendages,” extreme curvature of the spine (kyphosis and lordosis), especially elongated craniofacial bones, and large body size (Figure 1)(Dawson, 1985; Stiller et al., 2015). Substantial differences in these traits exist even among the three known extant species: *Phycodurus eques* (leafy seadragon), *Phyllopteryx taeniolatus* (weedy, or common, seadragon), and the recently described *Phyllopteryx dewysea* (ruby seadragon) (Stiller et al., 2015).

**Figure 1.**
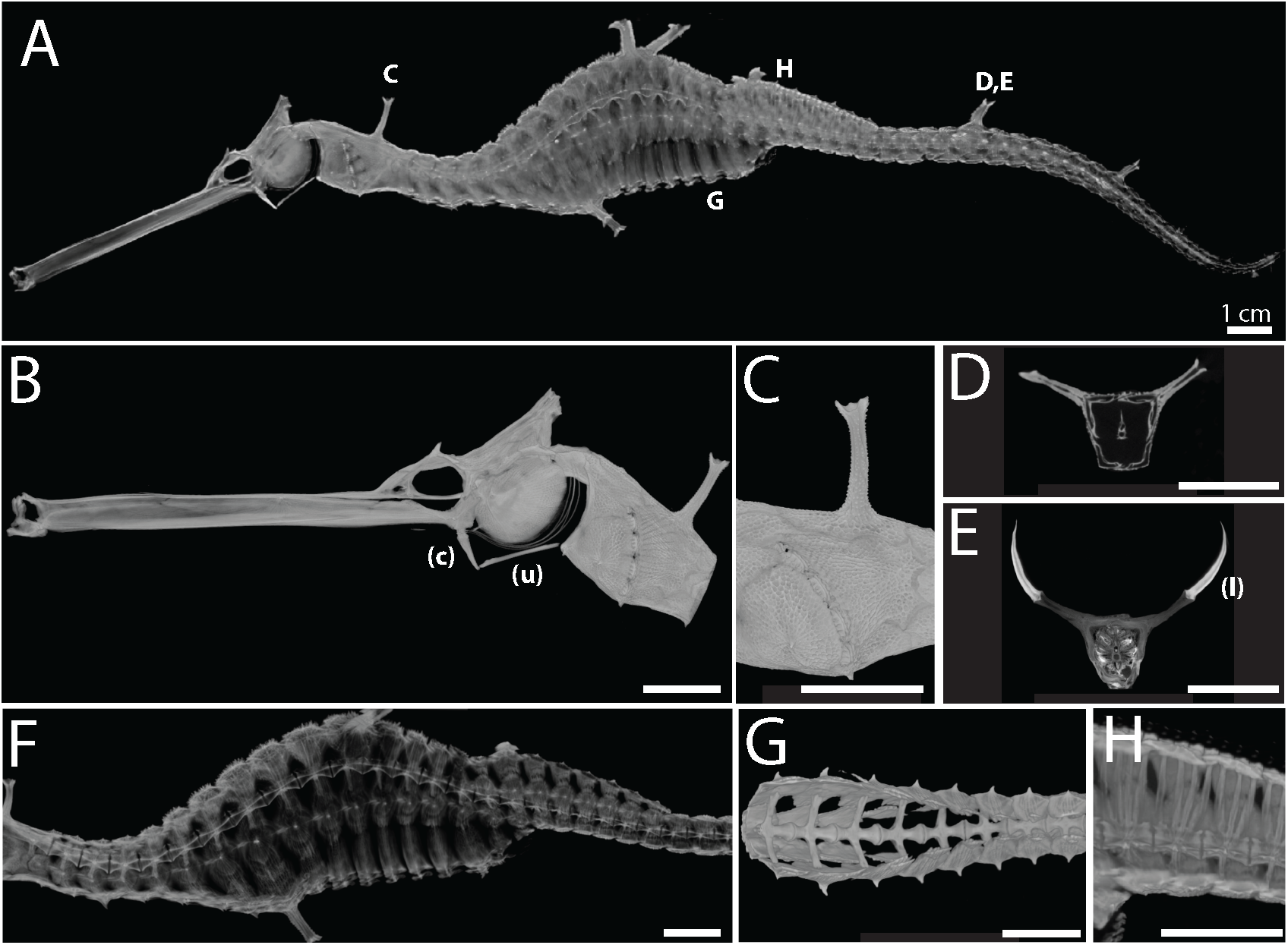
The anatomy of the weedy seadragon includes remarkably elongated facial features terminating in toothless, upturned jaws, an unusual hyoid apparatus specialized for suction feeding, a bony exoskeleton with elaborate spines that support fleshy “leaves”, and a sinusoidal spine of rib-less vertebrae that vary in shape and size. A) a lateral view of the skeleton of *Phyllopterix taeniolatus* reconstructed by X-ray microscopy. B) detail of the head (the ceratohyal, c, and urohyal, u, of the hyoid apparatus are noted). C) Detail of the pectoral region (lateral view) showing a dorsal, unpaired “leafy” appendage support surrounded by other dermal plates with much shorter spines. D) optical cross section of the tail through a pair of leafy appendage spines. E) optical cross section through the same appendages as in D) but with a contrast agent that reveals the fleshy leaves, seen edge on (denoted by l). F) lateral view shows keystone-shaped vertebrae at curvatures - both kyphosis and lordosis - of the spine. G) ventral view of the rib-less abdominal vertebrae. H) a lateral detail of the specialized vertebrae beneath the propulsive dorsal fin.

In addition to being a focus for evolutionary studies, seadragons are of significant cultural and conservation interest (Connolly et al., 2002; Martin-Smith & Vincent, 2006; Stiller et al., 2017). Presumed adaptations for crypsis including the leafy appendages, unique body plan, and elaborate skin coloration contribute to the status of seadragons as distinguished and valued cultural symbols for the people of Australia, where seadragon species are endemic. Because seadragon distributions are specific to temperate Australian macroalgal reefs, and their population sizes are relatively small, seadragons are likely susceptible to negative human impacts including global climate change. Furthermore, recent population genomic studies documenting significant population structure (Klanten et al., 2020; Stiller et al., 2021) in these species are especially relevant to conservation decisions. The unique evolutionary innovations, cultural importance, and conservation challenges all elevate the need to better understand and conserve seadragon species.

To improve our understanding of highly derived phenotypic traits and genomic features within seadragons, as well as those that are shared but derived among the Syngnathidae, we sequenced and annotated chromosome-scale assemblies for a male leafy seadragon and a female weedy seadragon. In addition to the production of these genomic resources we carried out several comparative analyses among five syngnathid and many other teleost genomes to determine changes in genome organization and content, including a more detailed analysis of a few key gene families and regulatory elements that may be involved in the development of syngnathid phenotypic innovations. Lastly, we performed high-resolution 3D X-ray microscope scans of an adult, male weedy seadragon to more precisely view seadragon morphological innovations.

Our work reveals a number of seadragon-specific genomic features, including divergent conserved non-coding elements (CNEs) near key developmental genes, a unique microRNA gene repertoire, and expanded gene families related to immunity and vesicular trafficking. We also found that the seadragon genomes are highly repetitive for their sizes, with unique repeat abundance distributions. Because the seadragon lineage occupies a region of the syngnathid phylogeny that is relatively basal to most of the species diversity, we leveraged their phylogenetic position to identify several genomic synampomorphies of the family. These genomic features include the striking loss of several highly conserved fibroblast growth factor (FGF) genes, expansions and contractions of gene families related to immunity and potentially male pregnancy, and syngnathid-specific transposable element (TE) expansion.

With these new genome models and rich accompanying data, we grow the existing collection of high-quality genomic tools and insights for several syngnathid groups including genera *Syngnathus* (Roth et al., 2020; Small et al., 2016), *Hippocampus* (Li et al., 2021; Lin et al., 2016), *Microphis* (Zhang et al., 2020), and most recently (published as of the writing of this paper) *Phyllopteryx* and *Syngnathoides* (Qu et al., 2021). We add to this quiver of syngnathid genomes useful in illuminating the evolution and development of puzzling syngnathid novelties such as male pregnancy and leaf-like appendages. These genomic resources will also support ongoing efforts to understand and conserve sensitive syngnathid populations, including phylogenetic umbrella species like the seadragons.

## MATERIALS AND METHODS

### Seadragon genome assemblies

We isolated high molecular weight genomic DNA from tissues of an adult male leafy seadragon (*Phycodurus eques*) and from an adult female common (“weedy”) seadragon (*Phyllopteryx taeniolatus*). We then generated PacBio libraries for both species and sequenced 49.12 and 80.80 Gb, respectively. We also generated “shotgun” whole-genome sequencing (WGS) Illumina libraries for both species, sequencing 57.48 and 105.79 Gb, to estimate genome size and polish the PacBio assemblies (see Supplemental Methods for all software versions and non-default parameters). We assembled both genomes with Flye (Kolmogorov et al., 2019), using all PacBio data excluding “scraps” with default parameters and an estimated genome size of 600 Mb. We performed two rounds of polishing on the primary Flye assemblies with the tool arrow (Chin et al., 2013), using the PacBio reads and performed an additional two rounds of polishing for each genome with WGS Illumina data, using pilon (Walker et al., 2014). To organize Flye assemblies into putative chromosome-scale genome models we generated Hi-C libraries using Phase Genomics Proximo Animal kits, then scaffolded using the 3D-DNA pipeline (Dudchenko et al., 2017) with breaking of original scaffolds disabled, followed by visualization and manual editing using Juicer and Juicebox (Dudchenko et al., 2018; Durand et al., 2016). We evaluated assembly quality and completeness using Quast (Gurevich et al., 2013) and BUSCO (Waterhouse et al., 2018).

### Draft short-read genome assemblies for *Doryrhamphus excisus* and *Synchiropus splendidus*

To supplement our comparative analyses with additional syngnathid genomes and a close outgroup we isolated high molecular weight DNA and generated linked-reads assemblies for the bluestripe pipefish (*Doryrhamphus excisus*) and the Mandarin dragonet (*Synchiropus splendidus*). These assemblies were performed using 10x Genomics Chromium technology and the Supernova assembly software (Weisenfeld et al., 2017). Details for these assembly methods are as described in Stervander and Cresko (2021).

### mRNA-seq

To generate mRNA-seq libraries, we extracted total RNA from tissues of the same *P. eques* and *P. taeniolatus* individuals as were used for PacBio genome sequencing. From the *P. eques* specimen, we dissected testis, leafy appendage, eye, and gill tissues. From *P. taenoiolatus* we dissected ovary, leafy appendage, eye, and liver tissue (see Supplemental Methods). We used the Roche KAPA HyperPrep Kit to generate indexed, stranded mRNA-seq libraries for Illumina sequencing to obtain 301.52 million paired-end 100 bp reads. We trimmed Illumina adaptors and low-quality regions from reads using process_shortreads from the Stacks software suite (Catchen et al., 2013; Catchen et al., 2011) and aligned cleaned RNA-seq reads from both seadragon species to both *P. taeniolatus* and *P. eques* genome assemblies using STAR aligner (Dobin et al., 2013).

### miRNA-seq

To generate small RNA-seq reads we purified a small fraction of RNA from each tissue above using Zymo DirectZol columns and generated indexed sequencing libraries using the NextFlex Small RNA-seq Kit v3. We ran BBMap (Bushnell, 2014) on the weedy and leafy seadragon genomes to align sequencing reads. We annotated miRNAs in the leafy seadragon based on sequence conservation with included species’ miRNAs (Antarctic blackfin icefish (*Chaenocephalus aceratus*), platyfish (*Xiphophorus maculatus*), European perch (*Perca fluviatilis*), and threespine stickleback (*Gasterosteus aculeatus*), zebrafish (*Danio rerio*), spotted gar (*Lepisosteus oculatus*), and medaka (*Oryzias latipes*)) as described in the Prost! (Desvignes et al., 2019) manual (see Supplemental Methods). Additionally, to run Prost!, we supplied threespine stickleback noncoding sequences (Desvignes et al., 2019). We ran Prost! for the weedy seadragon by adding the leafy seadragon’s annotations to the included species pool. We distinguished ohnologous miRNAs for the seadragon species using synolog (Catchen et al., 2009) synteny software, which aligned the leafy seadragon genome with the stickleback genome. Any miRNA not initially identified in the sequencing reads were searched for in the leafy seadragon genome with BLASTN and vista plots.

### Genome annotation

To facilitate the analysis of comparable annotations among genome assemblies, we performed new annotations for leafy seadragon (*P. eques*), weedy seadragon (*P. taeniolatus*), Greater pipefish (*Syngnathus acus*), bluestripe pipefish (*Doryrhamphus excisus*), Mandarin dragonet (*Synchiropus splendidus*), and Pacific bluefin tuna (Smit et al.) (*Thunnus orientalis*) (see Supplemental Methods for all public sequence data accessions). Assemblies were soft-masked for repetitive elements and areas of low complexity with RepeatMasker (Smit et al.) using custom repeat libraries made by combining a teleost-specific library extracted from RepeatModeler2 (Flynn et al., 2020) with species-specific repeat libraries produced by running RepeatModeler on each species listed above. We aligned all RNA-seq data (including new and previously published reads) as described above and supplied the .bam files to BRAKER2 (Bruna et al., 2021) for genome annotation.

We evaluated the final BRAKER2 set of annotated genes for each genome using InterProScan (Jones et al., 2014) and retained a final, “filtered” version of the annotation if they showed InterProScan evidence other than disorder-based (i.e. MobiDB-lite). We subjected remaining amino acid sequences to blastp searches against the NCBI RefSeq protein database and retained those in the filtered annotation if they returned a hit (e-value threshold of 0.001). For all downstream analyses of protein-coding sequences based on these and other annotations, we selected the longest transcript per locus.

### Repeat annotation

We characterized the repetitive content of 16 teleost genome assemblies (Figure S1; see Supp. Methods for accession numbers). With one exception (tiger tail seahorse (*Hippocampus comes*)), we focused exclusively on assemblies produced by long-read (i.e., PacBio or Oxford Nanopore) and/or linked-read (i.e., 10x Genomics) technologies. All genomes were subject to a unified repeat library generation and annotation workflow. Briefly, we identified repeats *de novo* for each assembly using 1. RepeatModeler2 (Flynn et al., 2020) and 2. TransposonPSI (Haas). We combined those repeat predictions with teleost repeats extracted from RepeatMasker (Smit et al.) libraries and all sequences from the FishTEDB database (Shao et al., 2018) http://www.fishtedb.org). These sequences (576,007 in total) were classified using the RepeatClassifier module of RepeatModeler. We also clustered repeats at 80% sequence identity using USEARCH (Edgar, 2010) as a way to group and ultimately enumerate repeats based on sequence divergence (see Supp. Methods). We ran RepeatMasker on all 16 genome assemblies using all 576,007 sequences and integrated the aforementioned RepeatClassifier and USEARCH cluster IDs into the RepeatMasker output. Finally, we used RepeatClassifier taxonomy and USEARCH cluster membership as alternative grouping mechanisms to characterize the distribution of repeats within and among genomes (e.g., using heatmaps and rank abundance plots), and to ordinate genomes in repeat space (i.e., using PCA). Regional repeat abundance distributions for target gene families were compared to those from random samples of genes across the genome using resampling-style hypothesis tests. All downstream repeat analysis and visualizations were carried out using the R statistical language (R_Core_Team, 2019).

### Gene family evolution

We used protein annotations from 21 teleost genomes (Figure 2; see Supp. Methods for accession numbers) to better understand gene family size evolution in seadragon and syngnathid lineages. We defined putative gene families via all-by-all blastp (Altschul et al., 1990) and clustering with mcl (Enright et al., 2002), then we conducted a series of gene family size evolution analyses using CAFE 5 (Mendes et al., 2020). Briefly, we first fit an error model to account for artifactual (e.g., genome assembly or annotation error) family size variation, which was applied to subsequent CAFE runs. We fit a model assuming a single rate of gene family expansion/contraction (λ) to the data and identified gene families evolving especially rapidly using CAFE’s internal likelihood ratio tests (LRTs).

**Figure 2.**
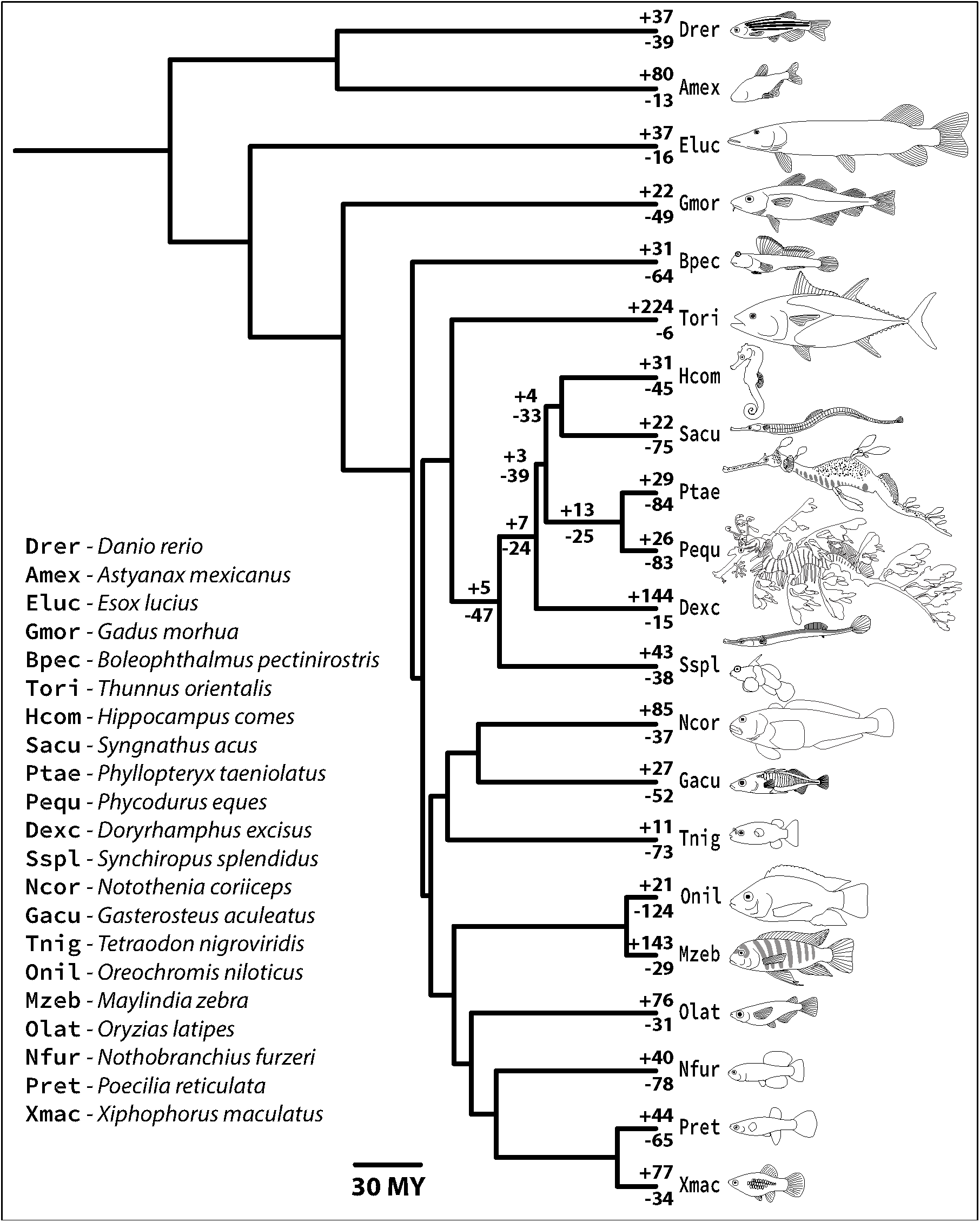
Genomes of 21 teleost species provide phylogenetic context for gene family evolution in seadragons and syngnathid fishes. Represented are evolutionary relationships among a sample of morphologically diverse teleosts, according to a time-calibrated phylogenetic tree adapted from Rabosky et al. (2018). We used published and newly generated protein-coding gene annotations for the species pictured here to understand putative gene family expansions and contractions in lineages of interest, specifically the seadragons. The number of gene families with statistical evidence for expansion (top values) and contraction (bottom values) along all terminal branches and internal syngnathiform branches are shown. Note that the four-letter species symbols in the legend are used throughout this article.

Of those gene families showing evidence for rapid evolution, we identified subsets for which branch-specific LRTs suggested extreme λs along branches of interest: the terminal *P. eques* and *P. taeniolatus* branches, the internal branch leading to the seadragon lineage, and the internal branch leading to the syngnathid lineage. We also assigned each gene family to a KEGG orthology (KO) ID if possible, and we used this information to perform KEGG pathway overrepresentation analysis using ClusterProfiler (Yu et al., 2012), with respect to the branch-specific rapidly evolving gene families. Lastly, we fit several multiple-λ models to the data in order to test hypotheses of overall λ differences (from a global λ) for the internal seadragon branch and the internal syngnathid branch. To test these hypotheses we used CAFE to perform 100 gene family simulations, fit the aforementioned models to the simulated data sets, and compared observed likelihood ratios (LRs) from the data to the LR distributions from the simulations. For details, see the Supplementary Methods.

### *fgf* and *fgfr* gene family characterization

We collected fibroblast growth factor (FGF) and receptor (FGFR) amino acid sequences from Ensembl of several percomorph species (see Supplementary Methods). We aligned orthologs and screened the genome assemblies of syngnathids and outgroups based on Hidden Markov Model (HMM) profiles from the alignments, according to methods described in Small et al. (Small et al., 2016). In some cases, we supplemented these sensitive searches with regional RNA-seq read alignments to correctly define exon boundaries. We conducted targeted, lineage-specific tests of positive selection using branch site-models in PAML (see Supplemental Methods) and tested for deleterious mutations using Provean (Choi & Chan, 2015).

### X-ray tomography and 3D model reconstruction

We obtained a euthanized adult male weedy seadragon, fixed in neutral buffered formalin from the Tennessee Aquarium (Chattanooga, TN), and scanned it using a resolution of 54 µm voxels on a Zeiss XRadia 620 Versa X-ray microscope at the University of Oregon Knight Campus for Accelerating Scientific Impact’s X-ray Imaging Core Facility. The anterior of the fish including the head and cleithrum were scanned again at 17 µm voxel resolution. Composite virtual three-dimensional reconstructions of the unstained specimen were generated using Dragonfly Pro and Dragonfly software (Object Research Systems). After completing scans of the unstained specimen, which allowed high contrast visualization of electron-dense bony structures, the fish was then dehydrated through an ethanol series, stained with an iodine-based X-ray contrast agent to enhance imaging of soft tissues, and a section of the rostral part of the stained tail that includes a pair of leafy appendages was scanned at 27 µm voxel resolution (Figure 1; Figure S2; Supplemental Methods).

## RESULTS

### PacBio assembly with Hi-C scaffolding yields chromosome-level genome models for the leafy and weedy seadragon

We estimated the haploid genome sizes for leafy and weedy seadragons to be 644.0 Mb and 597.3 Mb, respectively, based on k-mer frequency analysis of Illumina WGS data (Vurture et al., 2017). From this analysis we also estimated genome-wide heterozygosity for each individual to be 0.27 and 0.33%. Polished PacBio assemblies were 664.24 and 650.38 Mb, with scaffold N50s of 19.59 and 9.90 Mb. The longest Flye scaffolds were 38.02 and 29.83 Mb, and BUSCO completeness frequencies were 95% for both genomes. After scaffolding both polished PacBio assemblies using *P. eques* Hi-C reads, we obtained 23 putative chromosome models for each genome (Figure S3), which reflect 93.22% and 96.10% of the total length for final leafy and weedy seadragon genome models. These assembly and completeness metrics, our ability to annotate 22,256 and 22,043 protein-coding genes in the respective genomes, and extensive evidence for conserved synteny between the two seadragon assemblies (Figure S4), all support that these genome models are of high quality relative to available resources for teleost fishes. These genomes will therefore serve as useful tools for a variety of current comparative genomic analyses.

### Seadragon karyotypes are conserved relative to other syngnathid genomes but lack one of two chromosome fusions observed in *Syngnathus* and *Hippocampus*

Haploid chromosome number in syngnathid fishes, as assessed by karyotyping and genetic mapping, is reported to be 22 or 24 in seahorse (*Hippocampus*) species (Vitturi & Catalano, 1988; Vitturi et al., 1998) and 22 in Gulf pipefish (*Syngnathus scovelli*) (Small et al., 2016). In Gulf pipefish, the reduction in chromosome number from 24, the putative ancestral number in ray-finned fishes (Mank & Avise, 2006) to 22 likely resulted from fusion of two pairs of chromosomes orthologous to Chromosomes 1 and 24 and to Chromosomes 14 and 23 in platyfish (*X. maculatus*), a representative outgroup percomorph with 24 chromosomes (Small et al., 2016). Though a haploid number of 24 chromosomes in seahorse was reported (with some published confusion about in which species this was observed (Vitturi & Catalano, 1988; Vitturi et al., 1998) the reference genome for tiger tail seahorse (*H. comes*) (Lin et al., 2016), provides conserved gene synteny evidence that both ancestral chromosome fusions were already present in the common ancestor of seahorses and *Syngnathus* pipefish (Figure S5). As stated, our inferred seadragon haploid chromosome number of 23 is based on the size distribution of Hi-C scaffolds (Figure S3). The seadragon lineage apparently shares only the ancestral Chromosome 1 to Chromosome 24 fusion with the seahorse+*Syngnathus* pipefish ancestor, leaving one to one orthologs of platyfish Chromosomes 14 and 23 (Figure S5).

### Seadragon genomes are highly repetitive for their compact size, with large contributions from relatively recent TE expansions

A correlation between eukaryotic genome size and the proportion of a genome classifiable as TEs has led to an appreciation that TEs can be an important driver of genome size evolution (Kidwell, 2002). The relationship is particularly apparent among some of the well characterized genomes of teleost fish model species. Genome size and TE proportion share a positive, tightly linear relationship in a comparison of green spotted puffer (*Tetraodon nigroviridis*), threespine stickleback (*G. aculeatus*), medaka (*Oryzias latipes*), and zebrafish (*Brachydanio rerio*), whose genomes span a range from roughly 358 Mb to over 1.37 Gb (Gao et al., 2016). *Syngnathus* pipefish have genomes that rival green spotted puffer in genomic compactness, with assembly lengths of 307.0 Mb for the Gulf pipefish (Small et al., 2016) (*S. scovelli*), and 324.33 Mb for the greater pipefish (*S. acus*) (NCBI RefSeq Genome GCF_901709675.1). In the spectrum of vertebrate genomes, known seahorse genomes are also diminutive, estimated at 421 Mb for the lined seahorse (*H. erectus*) (Li et al., 2021) and 494 Mb for the tiger tail seahorse (*H. comes*) (Lin et al., 2016).

Our final genome assembly sizes for seadragons were 666.5 Mb for leafy seadragon and 652.2 Mb for weedy seadragon, which are appreciably larger than those of *Syngnathus* and *Hippocampus* but still much smaller than many other teleost genomes. Surprisingly, as much as 58.7% of the leafy and 57.9% of the weedy seadragon genome is composed of repetitive sequences, as classified by our workflow (see Methods). To assess whether seadragons or syngnathids are exceptional in their repetitive DNA characteristics, we measured the proportion of repetitive sequence in sixteen teleost genomes (Figure S1) using an in-common repeat reference library to create a standardized basis for comparison. Species were chosen for taxonomic breadth and because their genomes were assembled from long- or linked-read sequence data (except tiger tail seahorse). Genomes of five syngnathids were represented, including a seahorse and tail-brooding *Syngnathus* pipefish, the seadragons, and a basal lineage relative to these, the abdominal-brooding bluestripe pipefish (*D. excisus*).

We confirmed a strong, positive relationship between repeat content and genome size for the 16 genomes using phylogenetic generalized least squares (PGLS) regression (*t*_16,14_=4.88; *p*=0.0002; *b*=0.00048), but seadragons are notable outliers with large, positive residuals (Figure 3). Both seadragon genomes are unusually bloated with repetitive DNA among teleosts for their relatively compact size, a feature either derived specifically in the seadragon lineage or an ancestral lineage. This latter hypothesis is not yet testable given currently available genome assemblies but can be addressed with the addition of syngnathid genomes at key phylogenetic positions.

**Figure 3.**
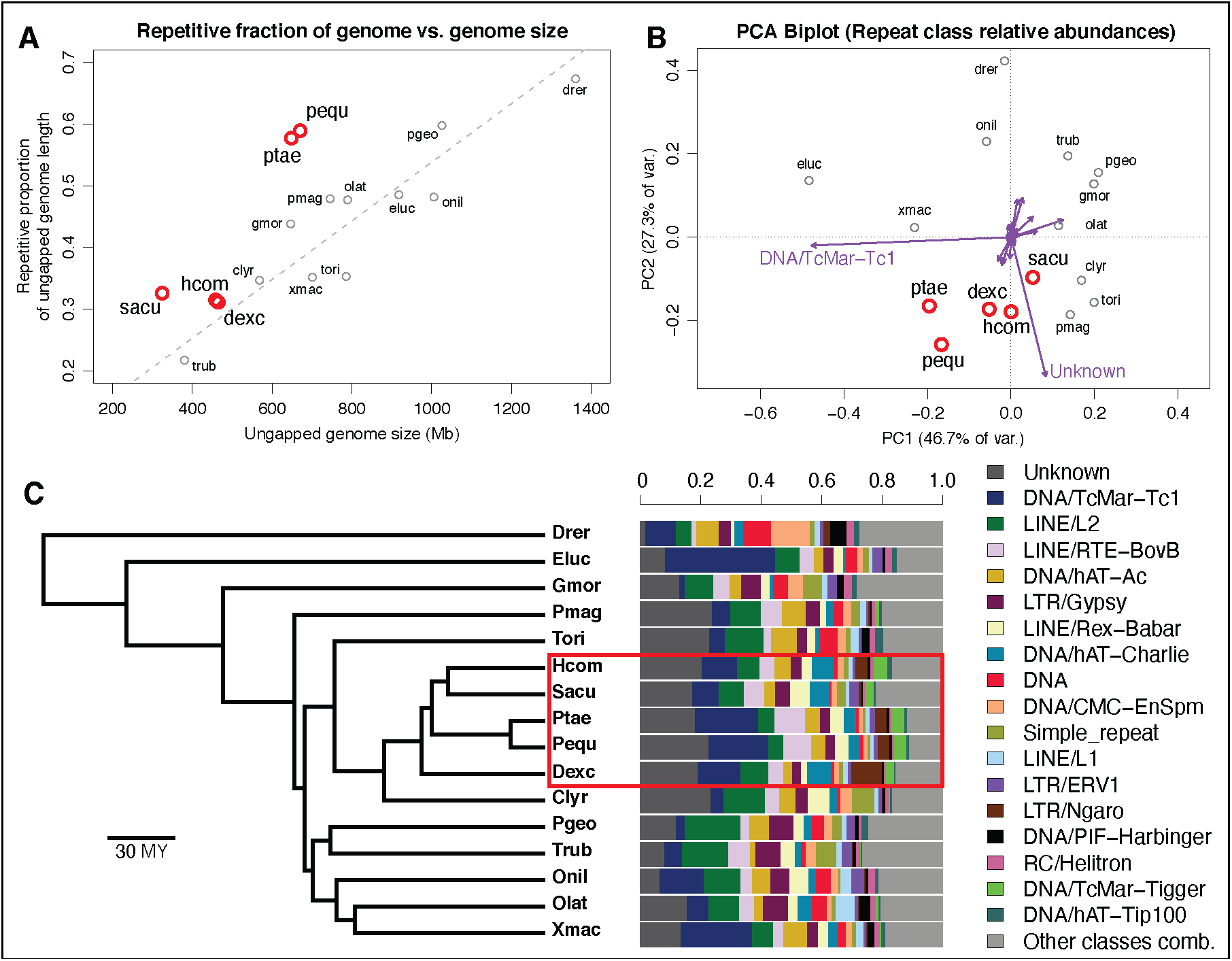
Seadragon genomes demonstrate a large fraction of repetitive DNA relative to other teleost fishes, characterized by substantial contributions from TcMar-Tc1 transposons and unclassified constituents. A) Scatterplot showing the strong, positive relationship between genome assembly size (x-axis) and the proportion of the genome annotated as repetitive for 16 recent teleost genome assemblies. Note that seadragon genomes (pequ and ptae) are especially repetitive (∼60%) given their relatively small size (∼650 Mb). Dashed line shows a phylogenetic generalized least squares (PGLS) regression fit, and syngnathid genomes are in red. B) A Principal Components Analysis (PCA) biplot shows similarity of the 16 genomes based on relative frequencies of repeat classes. TcMar-Tc1 and “Unknown” repeat classes load especially heavily on PC1 and PC2, respectively, as indicated by vectors (purple arrows) drawn in the space, and these contribute to the distinctiveness of syngnathid genomes (in red). C) Barplots showing the relative abundances of repeat classes across the 16 genomes, ordered (from left to right, and top to bottom in the legend) from highest to lowest mean relative abundance. Phylogenetic relationships among the 16 species are presented as a time-calibrated tree from Rabosky et al. (2018) and the syngnathid clade is indicated by a red rectangle.

Repeat density was high across seadragon chromosomes, in stark contrast with the greater pipefish (Figure 4), which shows repeat “hotspots” almost exclusively near chromosome ends, and a close outgroup to syngnathids, common dragonet (*Callionymus lyra*), which has relatively uniform, low repeat density in general (Figure S6).

**Figure 4.**
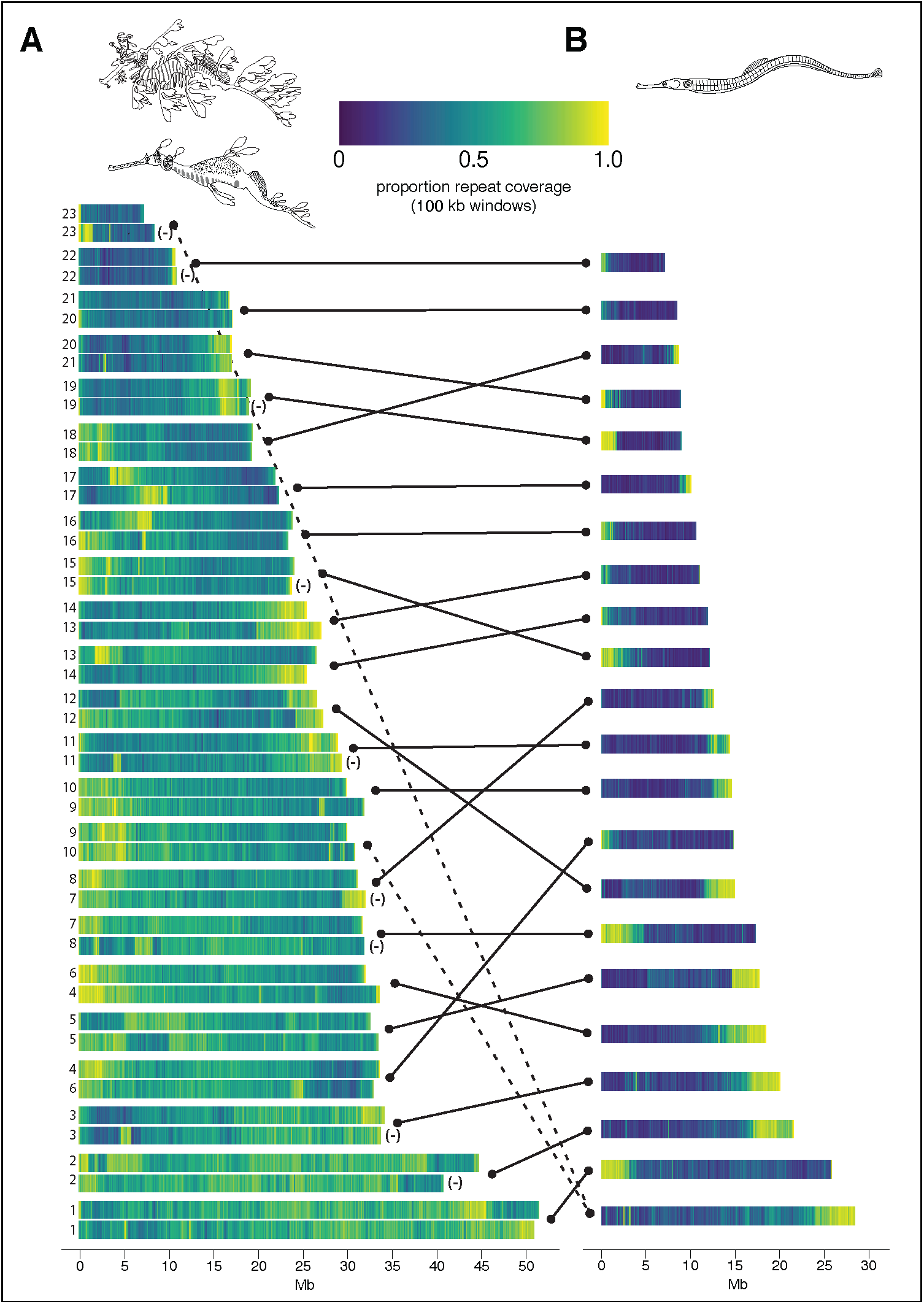
Leafy and weedy seadragon chromosomes are densely and non-uniformly populated by repetitive DNA. A) Orthologous pairs of the 23 seadragon chromosomes, ordered ascending from shortest to longest leafy seadragon sequence. For each pair the leafy seadragon chromosome is on top, and the weedy seadragon ortholog is below. Cases in which the orientation of the weedy seadragon chromosome has been reversed to align with the leafy ortholog are denoted by “(-)”. B) Chromosome models of the greater pipefish (*Syngnathus acus*), also ascending from shortest to longest. Lines connect seadragon and pipefish chromosomes with large regions of orthology, as inferred via conserved synteny analysis. Dashed lines reflect the fusion of two ancestral syngnathid chromosomes that is derived in the lineage leading to *Syngnathus* and *Hippocampus*. Overall repeat basepair occupation of 100 kb windows (expressed as a proportion) is presented as a heatmap.

Several TE classes contribute notably to the large repeatomes of seadragons. The *Tc1* family of the *Tc1/Mariner* superfamily of transposases is a major contributor to repeatome composition variation among teleosts as revealed by PCA based on within-repeatome relative class proportions, with seadragons, platyfish (*X. maculatus*), and Northern pike (*Esox lucius*) genomes influenced heavily by abundant *Tc1* repeats (Figure 3). Phylogenetic patterns of repeat abundance among the fish lineages we analyzed suggest *Tc1* expansion in the syngnathid lineage, given lower *Tc1* abundances in the close outgroups of common dragonet, (*C. lyra*), and Pacific bluefin tuna (*T. orientalis*). Among syngnathids, *Tc1* repeats compose a disproportionately large fraction of seadragon repeatomes (Figure 3). This class of “cut-and-paste” DNA transposons is widespread in animals and especially common in teleost fishes, with high abundance and variability among species (Gao et al., 2017; Gao et al., 2016). In fact, phylogenetic evidence suggests *Tc1* transposons are still active and recently expanding in some neoteleosts, such as threespine stickleback (*G. aculeatus*) (Gao et al., 2017).

The second most abundant, classifiable TE category in the seadragon genomes was the *BovB* family of non-LTR LINE retrotransposons (Figure 3; Figure S7). This TE family is restricted to animal taxa, where its members are patchily distributed among lineages and have been inferred via phylogenetic analysis to populate animal genomes through horizontal transfer, perhaps via metazoan parasites (Ivancevic & Chuong, 2020; Ivancevic et al., 2018). *BovB* density variation along seadragon chromosomes (Figure S7) was largely concordant with overall repeat density patterns (Figure S6), suggesting common mechanisms or constraints for the expansion of at least some TE families.

We also discovered an apparent expansion of *Tigger* transposases in the syngnathid clade, which are members of the *pogo* superfamily closely related to *Tc1/Mariner* (Gao et al., 2020). *Tigger* repeats are overrepresented in syngnathids compared to outgroups but proportionally are conserved among the five syngnathid genomes we analyzed (Figure 3). Unlike *Tc1* and *BovB, Tigger* repeat density variation along seadragon and *Syngnathus* pipefish chromosomes deviates from the collective pattern of repeat density (Figure S8), with the highest-density *Tigger* regions more centrally located, although the potential significance of this is unclear.

Because the grouping and enumeration of repeats according to classification alone comes with a loss of evolutionary resolution and precision, we also analyzed repeats as clusters of sequences with ≥ 80% sequence identity. PCA based on this cluster-wise treatment of repeats revealed that leafy and weedy seadragon genomes are quite divergent in repeat space from the other teleost species, including the three other syngnathids compared (Figure 5; Figure S9). In particular four repeat clusters are heavily influential in this regard, one of which was not classifiable but especially abundant in scaffolds unassigned to chromosomes (Cluster 20902), two of which belong to the *BovB* LINEs mentioned above (Clusters 10278 and 10626), and the last (Cluster 14395) belonging to the *Tc1* group of transposases mentioned above (Figure 5; Supplemental File 1).

**Figure 5.**
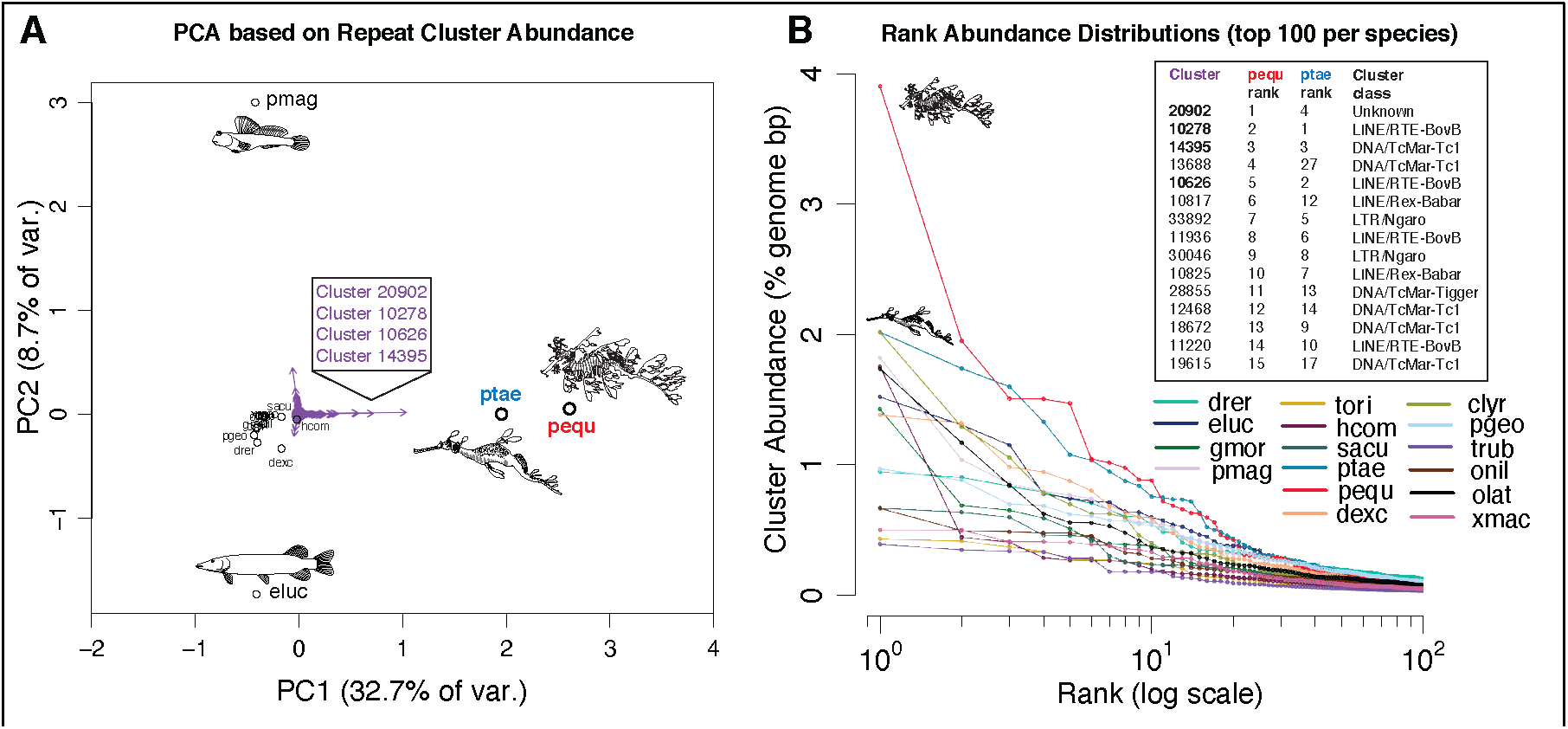
Relatively recent repeat expansions in the seadragon lineage drive the uniqueness of seadragon repeatomes. Repeat clusters defined at the 80% sequence identity level and quantified by the proportion of total genome length they occupy were used to conduct principal components analysis (PCA). A) A bi-plot of the first two PCA axes, showing extreme separation of seadragons from the other 14 species in repeatome space, particularly along the leading axis of variation (x-axis). Purple arrows represent individual repeat clusters, and how strongly (and in what direction) they influence the position of genomes in this repeatome space. Four repeat clusters (shown in a box) with especially large loadings on PC1 are strongly associated with seadragon repeatome uniqueness. B) Rank abundance distributions for the top 100 repeat clusters in each of the 16 species included in the repeat analysis. The top 20 clusters for each seadragon species are consistently elevated in abundance, relative to corresponding ranks in the other 14 fish genomes. Shown in a box are the top 15 leafy seadragon clusters, their ranks in weedy seadragon, and the repeat class to which they likely belong. Note that the top 3 (and 5^th^) clusters in the list correspond to the boxed clusters in panel A.

We also generated rank abundance distributions (RADs) for repeat clusters in the 16 fish genomes, an approach from ecology used to understand community evenness and major and rare constituents (Whittaker, 1965). Interestingly, leafy and weedy seadragon RADs show markedly high repeat cluster abundances relative to the other genomes for repeat ranks 1-20 (Figure 5). Among the highest-ranking repeats for both seadragon species are the four repeat clusters mentioned above as major contributors to the distinctiveness of seadragon repeatomes. These findings, along with the observation that members of the top-ranking seadragon repeat clusters are rare in the other genomes (Supplemental File 1), suggest that the unique repetitive features of seadragon genomes are driven largely by recent expansions of *BovB* and *Tc1*, and a yet-to-be-classified cluster.

### Gene family contractions in Syngnathidae involve innate immunity, and seadragon-specific expansions are associated with vesicular trafficking

We identified 290 total gene families as having expanded or contracted at a rate (λ) significantly higher than the background λ among 21 teleost species. Of these, 109 showed evidence for rapid size evolution along the leafy seadragon branch, 113 along the weedy seadragon branch, 38 along the internal seadragon branch, and 31 along the internal branch leading to syngnathids (Figure 2; Supplemental File 2). Based on 100 simulations of gene family evolution along the tree, we also inferred that λs for the internal seadragon and syngnathid branches are respectively distinct from the global λ for the tree, and likely distinct from one another (Figure S10).

Among the gene families most likely to have evolved rapidly in size in the ancestral syngnathid lineage, at least seven families related to innate immunity experienced contractions (Supplemental File 2), in contrast to multiple lines of evidence for expansion of inflammation and innate immunity gene families in teleosts relative to other vertebrates (Balla et al., 2020; Boudinot et al., 2011; Chang et al., 2021; Howe et al., 2016; Mattingsdal et al., 2018). Specifically, we found evidence for syngnathid-specific contractions in NACHT, TRIM, and GIMAP gene families, consistent with some of the immunity and detoxification pathway gene families depleted in the genome of the Manado pipefish *(Microphis manadensis*) (Zhang et al., 2020).

Several functional categories (KEGG pathways) were overrepresented among gene families with large size changes in the seadragon lineage, including cancer, cardiomyopathy, and immunity (Supplemental File 2), primarily due to contraction events. In terms of gene family expansions along the branch leading to seadragons, however, two families with roles in vesicular trafficking - *Vacuolar Protein Sorting-Associated Protein 13B* (*vps13b*) and *Coatomer Protein Complex Subunit Beta 2* (*copb2*) – are notable. Sequences with high similarity to *copb2*, which encodes one subunit of a Golgi budding and vesicular trafficking protein complex (Waters et al., 1991), are especially abundant in seadragon genomes relative to other syngnathids and teleosts (Supplemental File 2).

Though tuna (*T. orientalis*) and platyfish (*X. latipes*), for example, have two paralogs of *copb2*, we could find evidence for only a single gene copy in the genomes of seahorse (*H. comes*) and pipefish (*S. acus*). By contrast, we detected the presence of at least nine and six copies (on the 26 longest Hi-C scaffolds) in leafy and weedy seadragon genomes (Supplemental File 3). We also found many sequences matching *copb2* on several short scaffolds, suggesting that the repetitive nature of this region prevented these from being incorporated into chromosome-scale scaffolds, or, but less likely (Figure S11), that they could be redundant artifacts. One of the *copb2* paralogs (on Pequ Hi-C scaffold 13, Ptae Hi-C scaffold 14) is the likely ortholog of platyfish *copb2* on the orthologous chromosome (Xmac 6), with the remaining copies likely expanding secondarily via an unknown mechanism.

Given the repetitive nature of these regions, we hypothesized that TE activity may have played a role in seadragon *copb2* expansion. Specifically, we first tested whether the seadragon-expanded TE classes of *BovB* and *Tc1* are overrepresented in the immediate vicinity (1-kb flanking both sides) of *copb2* copies, relative to 499 randomly resampled gene groups. Both *BovB* (4.24-fold enriched; *p* < 0.002) and *Tc1* (1.39-fold enriched; *p* < 0.002) repeat classes were significantly overrepresented in *copb2* regions (Figure S11; Supplemental File 4). We secondarily performed naïve hypothesis tests with the false discovery rate (FDR) controlled at 0.05 to identify other TE classes with potential enrichment in these regions, revealing *Rex-Babar* LINEs, and *Tigger* and *Charlie* transposons as additional candidates (Supplemental File 4).

### Syngnathid fishes have lost several FGF family genes, most notably *fgf3* and *fgf4*

The FGF and FGFR gene families include well-studied ligand and receptor signaling molecules central to vertebrate craniofacial, limb, dermal appendage, sensory placode, and hindbrain development, among many other functions (reviewed in Xie et al., 2020). Because of the prominence of FGF signaling in the morphogenesis of traits that are distinctively modified in syngnathids, we explored whether FGF ligands or their receptors are exceptional in seadragons and in three other representative lineages of syngnathids: flagtail pipefishes (the most basal lineage of the four), seahorses, and *Syngnathus* pipefishes (Figure 2).

The FGF and FGFR families in seadragons and other syngnathid fishes conform, with two notable exceptions mentioned below, to a broadly typical complement of gene paralogs relative to other percomorphs. Within the Syngnathidae, there are several lineage-specific losses of FGF and FGFR genes (Table S1). We did not detect *fgf9* in any syngnathid. Seadragons and *Syngnathus* pipefish have apparently lost *fgf4-like*,

while tiger tail seahorse (*Hippocampus comes*) and bluestripe pipefish (*Doryrhamphus excisus*) have retained it. *fgf17* is missing birds (Abramyan, 2015), in seadragons and bluestripe pipefish but is present in tiger tail seahorse and greater pipefish. Because flagtail pipefishes, the lineage to which bluestripe pipefish belongs, are basal to the clade containing seadragons, seahorses, and *Syngnathus* pipefish (Longo et al., 2017), *fgf17* therefore must have been lost at least twice. Evolutionary loss of *fgf17*, though uncommon, is not unique to syngnathid lineages; orthologs of this gene have been independently erased from other distant taxa, such as the medaka genus *Oryzias* (Canestro et al., 2007). In addition, at least two clades of *fgf5* and *fgfr1b* are present in seadragons, tiger tail seahorse, and blue-stripe pipefish, but these genes were not detected in the two *Syngnathus* pipefish genomes (*S. acus* and *S. scovelli*).

The most startling gene losses from the FGF family are shared across the seadragon, tiger tail seahorse, greater pipefish, and bluestripe pipefish lineages: all are missing *fgf3* and *fgf4* (Figure 6; Table S1). In percomorph outgroups to the syngnathids, *fgf3, fgf4*, and *fgf19* are clustered, with *fgf3* flanked by *ccnd1* and *lto1*, genes that are also missing from seadragons, seahorses, and greater pipefish (but bluestripe pipefish has retained *lto1* (Figure 6). The expected neighbors normally on the *fgf19* side of the cluster (*zgc:153993* and *ano1*) are present in seadragons and both lineages of pipefishes, but bluestripe pipefish has lost *fgf19*. Close outgroups to the syngnathids, the razorfish (*Aeoliscus strigatus*) and the mandarinfish (*Synchiropus splendidus*) (Longo et al., 2017) retain intact *fgf3/4/19* clusters, making it likely that drastic alteration to the cluster is a syngnathid synapomorphy (Figure S12). In the common ancestor of lobe-finned and ray-finned fishes, *fgf3, fgf4*, and *fgf19* were likely already clustered, and remain tandemly arrayed in representatives from jawed vertebrate lineages as separated as mammals and teleosts (Oulion et al., 2012). The syngnathid lineage has therefore experienced a degree of change in this cluster that is perhaps unprecedented throughout gnathostome evolution.

**Figure 6.**
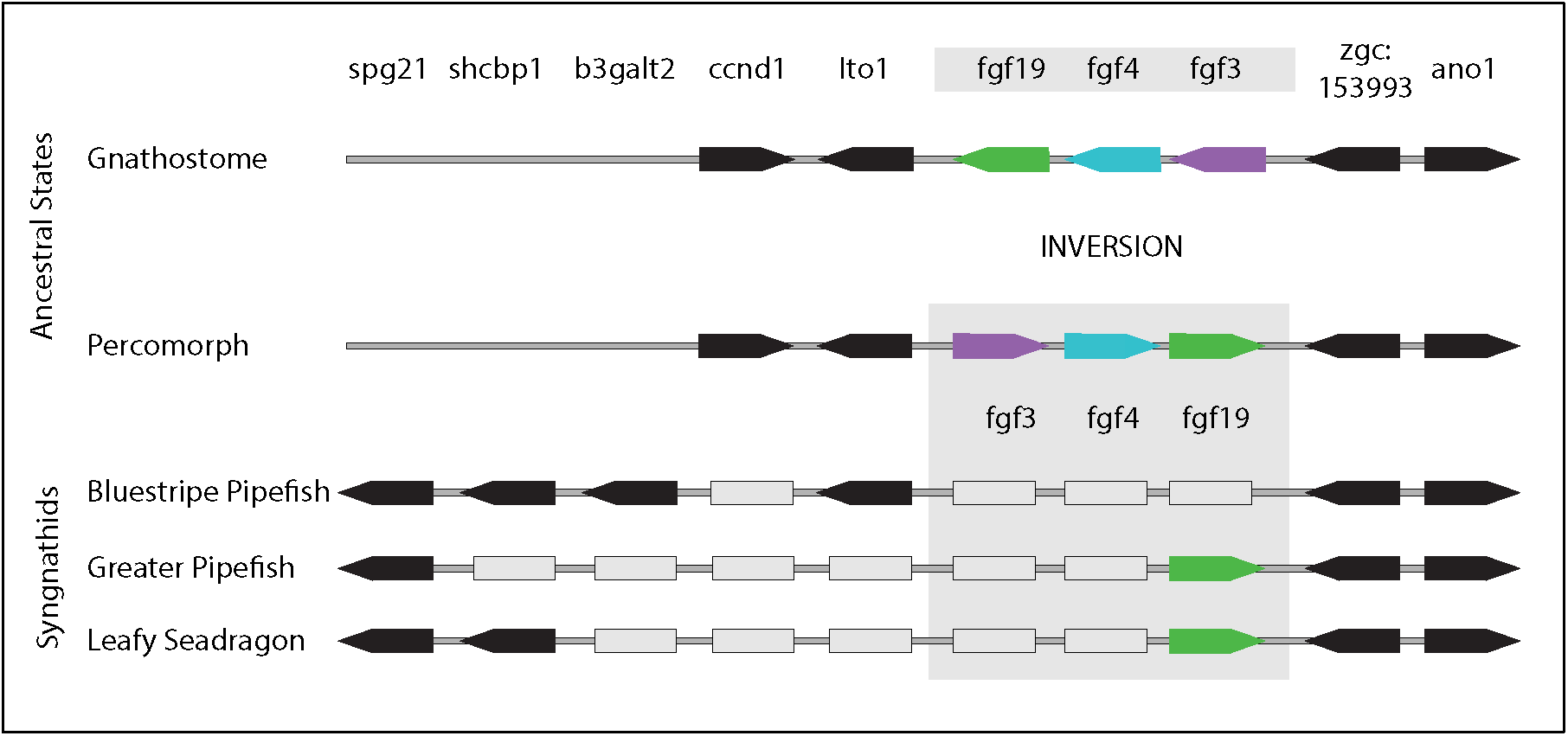
The *fgf3/4/19* cluster locus has experienced surprising gene losses in the syngnathid lineage. While retaining the same immediate gene neighbors, the FGF cluster became inverted in the percomorph fishes relative to outgroups like zebrafish and tetrapods. Several genes (grey rectangles) appear to have been deleted from the locus in the syngnathids, though not all of the losses are shared. *spg21, shcbp1*, and *b3galt2*, which neighbor the locus in the syngnathid lineage, are separated from the FGF cluster by other genes in non-syngnathid percomorphs like platyfish. Genes that appear lost from the locus cannot be found anywhere else in the genome assemblies of pipefishes, seahorses or seadragons. Arrows show gene order and orientation; grey rectangles represent apparent gene losses.

Exceptional absences of FGF and FGFR members in extant syngnathid genomes suggest that some property of their syntenic neighborhoods is inherently volatile. We tested whether the local repeat landscape flanking significant regions of FGF, FGFR, and other developmental gene loss in syngnathids – the *fgf3/4/19* cluster, *fgf4l, fgf5, fgf17, fgfr1b, eve1* (Small et al., 2016), and *tbx4* (Lin et al., 2016; Small et al., 2016) - differs from that of genes in general, by using the leafy seadragon genome and the close syngnathid outgroup *C. lyra* (common dragonet) as a pre-loss comparator. While no individual repeat class was significantly enriched in these regions (of either genome) after FDR adjustment, several show nominal evidence for enrichment and are at high density in the leafy seadragon regions, including *Tc1, BovB, hAT-Ac*, and additional LINEs *Rex-Babar*, and *L2* (Figure S13; Supplemental File 4). Furthermore, we found support for the alternative hypothesis that *overall* repeat density in general is enriched in these regions (Figure S13; Supplemental File 4), but only for the leafy seadragon (Pequ: 1.45-fold enrichment; *p* = 0.022), and not the dragonet genome (Clyr: 1.11-fold enrichment; *p* = 0.27).

### Limited evidence for compensatory evolution in the FGF pathway

FGF paralogs sometimes overlap in their expression and ability to compensate, partially or fully, for loss-of-function of one another in specific developmental contexts. In zebrafish, for example, knock down of either *fgf3* or *fgf10* causes relatively mild effects in lateral line migration and neuromast maturation, but there are severe lateral line defects when expression of both paralogs is depleted (Nechiporuk & Raible, 2008). We explored whether those paralogs most recently diverged from *fgf3* and *fgf4*, or with overlapping developmental roles, showed evidence for compensatory evolution prior to the syngnathid radiation.

Syngnathid and close outgroup sequences are largely conserved for *fgf4-like, fgf6a, fgf7, fgf8a, fgf8b, fgf10a, fgf10b, fgf19, fgf20a, fgf20b, fgf22*, and *fgf24*, and exhibit no evidence of positive selection along the syngnathid branch (Table S2). We did find potential evidence for positive selection in *fgf6b*, although this test result was within the range of false discovery (Figure S14; Table S2). Second, although the gene was lost in the seadragons and blue striped pipefish lineages, we found the strongest evidence for positive selection on *fgf17*, which is retained in seahorse and *Syngnathus* pipefish (Table S2).

Like the ligands, the syngnathid FGF receptors are largely conserved, and we found no evidence of syngnathid lineage-specific positive selection for non-canonical *fgfrl1a* and *fgfrl1b*, nor for canonical *fgfr1a, fgfr2, fgfr3*, or *fgfr4*. In contrast, *fgfr1b* presented some evidence for positive selection (although not robust to false discovery), displaying surprising syngnathid-specific substitutions (Table S2). These include a likely deleterious substitution within the activation loop of the kinase domain, a position conserved not only across vertebrate Fgfr1 proteins, but across all canonical FGFR paralogy groups (Figure S15) (provean score= -6.123).

Although we did not find evidence for lineage-specific positive selection using codon-based models, leafy and weedy seadragons share a derived six amino acid deletion in a conserved region of the Fgf16 protein (Figure S16), while seahorse has amino acid substitutions in this same motif. Other representative percomorphs, chicken, and human Fgf16 protein sequences are identical across this region indicating a high level of conservation, though no function has yet been ascribed to this protein domain. Divergence of seadragon *fgf16* is not limited to the coding sequence. A putative regulatory change is hinted at by the absence in both dragons of an approximately 240 bp conserved non-coding element (CNE) that is well preserved across percomorph fishes 5’ of *fgf16* (Figure S17).

### miRNA-seq data from seadragons reveal loss of conserved microRNAs

Although microRNAs (miRNAs) are important developmental regulators (Bizuayehu & Babiak, 2014), there are currently no annotations of these genes in syngnathids. In the leafy and weedy seadragons, we identified 261 and 251 miRNA genes that produce 331 and 318 unique mature miRNAs (Supplemental File 5). We found absences of numerous conserved miRNA genes including *mir10a* and *mir196b*, hox miRNAs genes that were reported missing in the Gulf pipefish genome assembly (Small et al., 2016). Six of the additional highly conserved missing miRNAs belong to two miRNA clusters from the miR-130 family. These absences include *mir130a, mir301a, mir130b, mir301b, mir130c-2*, and *mir454b*. While *mir130a, mir301a, and mir301b* are convergently missing in platyfish (*X. maculatus*) and medaka (*O. latipes*), absences of *mir130b, mir130c-2*, and *mir454b* have not yet been reported in other vertebrate species (Figure S18) (Desvignes et al., 2019; Desvignes et al., 2021; Kelley et al., 2021). In some tetrapods and teleosts, *mir301a* is located in the first intron of *ska2*. The seadragons, Gulf and greater pipefishes, and tiger tail seahorse are missing *ska2* as well as *mir301a* and *mir130a*. Throughout vertebrates, *mir130b* and *mir301b* (and, in teleosts and Coelacanth, additionally *mir130c-2* and *mir454b*) are located in the intergenic region between *sdf2l1* and *top3b*. Though these adjacent protein coding genes and the immediate syntenic neighborhood are conserved in seadragons, Gulf and greater pipefishes, and tiger tail seahorse, this cluster of microRNAs is not.

## DISCUSSION

Exploring the seadragon genomes in a comparative phylogenetic context has lifted a veil on the evolution of seadragon-specific traits and has also revealed intriguing evolutionary facets of this unusual vertebrate family, the Syngnathidae, as a whole. We found that both leafy and weedy seadragon genomes stand out among their relatives in having a surprisingly large contingent of repetitive DNA. This pattern appears to have been driven largely by recent expansions of *BovB* and *Tc1*, and a yet-to-be-classified cluster of repetitive sequences specific to the seadragon lineage. One possible explanation for the large difference we observed between seadragon and *Syngnathus* pipefish repeatome size and genomic distribution could be a difference in historical effective population size (N_e_). In this case, negative selection to remove rapidly expanding repeats from a population would be less effective in the face of strong genetic drift (small N_e_) (Lynch, 2007; Lynch & Conery, 2003). Differences in k-mer-based heterozygosity estimates from individual WGS data are at least consistent with this idea. A heterozygosity estimate based on the Gulf pipefish data (Small et al., 2016) is roughly three times the seadragon estimates (1.01% versus 0.27% and 0.33% for leafy and weedy seadragons, respectively).

The explanation presented above for TE expansion assumes that deleterious effects are common, likely through interruption of coding regions or promoters, or by regional silencing via chromatin changes (Liu et al., 2018). However, TEs can also contribute new genes or gene regulatory sequences when a host genome co-opts (“domesticates”) these exogenous genetic elements (Bejerano et al., 2006). Some classes of TE provide, for example, binding sites for master transcriptional regulators NANOG and OCT4 throughout the mouse and human genomes but in largely non-overlapping sets of loci between the taxa, potentially restructuring, in lineage-specific ways, the transcriptional networks for developmental pluripotency (Kunarso et al., 2010). TEs and repetitive DNA in general can fuel gene family expansion or contraction by precipitating unequal crossover events; Hahn et al. (2007) suggest an “explosion” of TEs in the primate lineage could be linked to its accelerated gene content evolution in primates. Repeatome expansion, such as what we here observe in seadragons, could have had disruptive impacts on gene regulatory networks and gene content, subject to subsequent evolution via negative *or* positive selection.

Using global and targeted approaches we explored expansion and contraction of gene families in seadragons and their relatives, and possible connections of gene content changes to observed TE distributions. Perhaps the most obvious trend we observed for the syngnathid lineage in general was contraction of particular immunity and detoxification pathway gene families, some of which have previously been described (Lin et al., 2016; Zhang et al., 2020). GTPase of the immunity-associated protein genes (GIMAP), for example, was among the contracted families in seadragons that was also detected in the seahorse genome (Lin et al., 2016), and our more comprehensive analysis supports that the GIMAP contraction occurred prior to the radiation of syngnathids. This family exists as a single eight-member cluster in mammals (Krucken et al., 2004). GIMAP genes have multiplied in other teleost lineages, numbering up to nearly 190 genes. Balla et al. (2020) found zebrafish GIMAP genes respond to pathogenic viral exposure and suggest the gene expansion could have been evolutionarily favored by the relatively long period that hatchlings must rely on innate immunity before the development of a functional adaptive immune system. Male-brooded syngnathid embryos might enjoy a luxury not available to free-spawned progeny like those of zebrafish, namely pathogen climate control afforded by the paternal immune system before their own adaptive immunity develops. The male’s immune system strikes a balance of defending against foreign agents but without rejecting his own brood. The syngnathid contraction of the GIMAP family is also interesting, therefore, in light of the possibility that *gimap4*, which does persist in the lineage, could promote immunologic tolerance to embryos in brood pouch tissues (Roth et al., 2020). Roth et al. (2020) based this assertion on the observation that *gimap4* is up-regulated in *Syngnathus* pregnancy tissues, where it could contribute to local suppression of the lymphocyte population (Roth et al., 2020; Schnell et al., 2006).

We detected seadragon-specific copy number expansion of a coatamer complex gene *copb2*. Variants of the *BovB* and *Tc1* transposable elements were enriched surrounding the supernumerary gene copies, suggesting a TE-driven mechanism for the expansion. Copb2 forms part of a protein complex involved in retrograde vesicle budding from the Golgi apparatus and secretion of macromolecule cargo, such as collagen, which is critical for bone and connective tissue development. Mice and fish developing with deficits in *copb2* have delayed bone mineralization and low bone density, as well as defects in type II collagen trafficking and secretion (Marom et al., 2021). Zebrafish mutants of *copb2* develop mispatterned, kinked notochords with poorly formed vacuoles and a disorganized perinotochordal basement membrane, which is secreted by notochord sheath cells (Coutinho et al., 2004). In teleosts, the notochord, particularly its sheath, plays an instructive role in patterning the vertebrae (Gray et al., 2014; Peskin et al., 2020).

Given the elaborated bony exoskeleton in seadragons, their stiff bodies with connective tissue-dense leafy ornaments, and their kinked axial skeletons with varied and regionalized vertebral forms (Figure 1), the proliferation of *copb2* gene sequences in their genomes ignites curiosity about the possible evolutionary developmental consequences of this expansion for the seadragons’ unique exo- and endoskeletons. Kinked vertebral columns in the guppy (*Poecelia reticulata*) mutant *curveback* are characterized by wedge-shaped vertebrae (Gorman et al., 2007). Similar to the guppy mutant and to the developmental malformation of vertebrae in human Scheuermann’s kyphosis sufferers, vertebrae are keystone-shaped in the weedy seadragon at locations of spinal curvature (Figure 1; Figure S2).

One of the most surprising gene family reductions we uncovered is shared by all of the syngnathid lineages we explored; it is the loss of *fgf3* and *fgf4*. Loss of these two fibroblast growth factor genes in syngnathids is striking because their orthologs in other vertebrates have long been hypothesized to play nearly indispensable pleiotropic developmental roles in the pharyngeal arches, teeth, brain, cranial placodes, epidermal appendages, limbs, and the segmental axis (Anderson et al., 2020; Boulet et al., 2004; Cooper et al., 2017; Cooper et al., 2018; Crump et al., 2004; David et al., 2002; Jackman et al., 2013; Leger & Brand, 2002; Lu et al., 2006; Maves et al., 2002; Miyake & Itoh, 2013; Nechiporuk & Raible, 2008; Prykhozhij & Neumann, 2008; Reuter et al., 2019; Walshe et al., 2002; Wang et al., 2007). It is reasonable to weigh whether losing these two multifunctional signaling ligands could have had significant and broad consequences to both deeply conserved developmental pathways and their morphological readouts. Another possibility is that these developmental pathways had diverged neutrally, or through changes in other pathway members, from anciently conserved functions along the syngnathid lineage, permitting genes that had once been critical to become expendable. It is nevertheless valuable to note that syngnathid fishes share peculiarities in many features from the constellation of vertebrate traits that *fgf3* and *fgf4* are known to help pattern.

Dermal integuments of syngnathid fishes are bony plates, and in several syngnathid lineages including the seadragons these have been elaborated to magnificence, sometimes independently. A subset of these plates in seadragons bear blunted struts of bone that end in fleshy paddle-shaped ornaments, the “leaves” and “weeds” of the species featured in this report (Figure 1). Another percomorph clade, the pufferfishes, are adorned with bony spines likely evolved from elasmoid scales. Shono et al., (Shono et al., 2019), showed that *fgf3* (and *fgf1a*) are expressed in developing pufferfish dermal spines. Given this and a trove of other evidence for an FGF signaling role in the development and diversification of scales, spines, and denticles in ray-finned and chondrichthyan fishes (Albertson et al., 2018; Aman et al., 2018; Cooper et al., 2017; Cooper et al., 2018; Daane et al., 2016; Kim et al., 2019; Shono et al., 2019), absence of *fgf3* and *fgf4* in the often heavily armored, elaborately spined syngnathids clearly suggests that derived mechanisms for integumentary bone development are at play in this lineage.

Syngnathids have evolved elongated faces with an unusual hyoid apparatus integral to specialized suction feeding (Figure 1) (Leysen et al., 2010), and they are toothless. These craniofacial features arise developmentally from complex interactions between - at least - endoderm, mesoderm, and neural crest components. Both *fgf3* and *fgf4* are expressed in the pharyngeal arches, which form skeletal elements of the jaw, hyoid, and, in fish, the gill supports (David et al., 2002; Herzog et al., 2004; Niswander & Martin, 1992) *fgf3* plays an essential role in craniofacial development. When *fgf3* is knocked out in zebrafish, mutants die within 7-9 days; craniofacial defects (particularly the full loss of cartilage in gill arches) are predicted to be the cause of their early mortality (Herzog et al., 2004). Additionally, when *fgf3* expression is disrupted in the mesendoderm in developing zebrafish, an “inverted” backward directed ceratohyal cartilage is formed (David et al., 2002). It is tempting to speculate that evolutionary loss of these genes could have led to altered craniofacial architecture of the elongated syngnathid face, either directly or through the effects of genetic compensation percolating through this signaling pathway.

Syngnathid toothlessness is also particularly interesting given *fgf3* and *fgf4* expression in zebrafish dental epithelium and their suspected roles in tooth morphogenesis (Jackman et al., 2004). Gilbert et al. (2019) propose a model for patterning of dentition in ray-finned fishes in which a tooth primordium acts as an organizer that induces the development of subsequent teeth, likely via secretion of Fgf3 and Fgf4. Other tooth developmental genes are known to be lost or reduced in copy number in syngnathids, including *eve1* (Small et al., 2016); see above) and P/Q-rich SCPP enamel/enameloid matrix genes (Lin et al., 2016; Qu et al., 2021; Zhang et al., 2020). These losses imply an erosion of tooth development pathways spanning induction to mineralization, with our discovery of *fgf3/4* loss enlarging the pool of candidate causative genes for edentulism in syngnathids.

The syngnathid central nervous system features its own peculiarities. Bennedetti (1991) described greater pipefish (*S. acus*) and long-snouted seahorse (*Hippocampus guttulatus*) to have highly modified or no discernable Mauthner neurons, the large, rhombomere 4 (r4) reticulospinal neurons critical for the rapid “C-start” escape response in many fishes and some amphibians (reviewed in Korn & Faber, 2005). Syngnathids are reputed also to lack mechanosensory lateral line neuromasts, from which the Mauthner cells receive synaptic inputs (Benedetti, 1991; Korn & Faber, 1975). In zebrafish, joint impairment of *fgf3* and *fgf8* impacts segmental identity of rhombomeres 3 and 5 and their reticulospinal neurons (Maves et al., 2002; Walshe et al., 2002). Furthermore, depletion of *fgf3* and *fgf10* reduced the number of zebrafish posterior lateral line neuromasts and inhibited migration of the placode down the length of the body (Nechiporuk & Raible, 2008). These observations present a compelling case for future interrogation of a link between *fgf3* loss and derived hindbrain and sensory development in syngnathids.

Though it is known that paralogous FGF ligands can compensate for one another in some experimental contexts (Maves et al., 2002; Walshe et al., 2002), in general, we did not find sweeping evidence for adaptive evolution of paralogous FGF proteins or their receptors in the wake of syngnathid *fgf3* and *fgf4* gene losses. Correlated changes could instead have included evolution of non-coding, regulatory sequences of FGF/FGFR genes or changes to other gene families that interact with FGF signaling. Syngnathids, we found, have lost six deeply conserved miRNAs in the miR-130 family. Biological implications for these missing miRNA genes are uncertain, though it is possible that their losses could relate to derived syngnathid-specific traits and gene pathway changes. For instance, angiogenesis and tissue remodeling are critical components in syngnathid male pregnancy tissues (Stolting & Wilson, 2007), and mir130a is connected to vascular repatterning in mammals and teleosts (Chen & Gorski, 2008; Singh et al., 2017) mir130 genes are also known to affect FGF signaling. In chicken, knockdowns of *mir130* lead to increased expression of *fgf8* and FGF pathway member, *Erk 1/2* (Bobbs et al., 2012; Lopez-Sanchez et al., 2015). There is also a suspected regulatory relationship between FGF receptors and these miRNAs; in human differentiating stem cells, nuclear located FGFR1, nFGFR1, binds to the promoters of *mir301b and mir130b* (Stachowiak & Stachowiak, 2016; Terranova et al., 2015). Loss of miRNAs that regulate and are potentially regulated by FGFs and FGFRs could indicate a further extension of the derived restructuring of FGF signaling pathways in syngnathids.

Evidence for positive selection as a compensatory consequence of having lost *fgf3* and *fgf4* in a syngnathid ancestor proved to be scarce. *fgf16*, however, provides a possible example of a separate evolutionary scenario: derived functional change in the seadragon lineage, at both the protein and cis-regulatory levels. Leafy and weedy seadragon fgf16 proteins share an unusual deletion in a deeply conserved motif, and the seadragon gene also appears to have lost a 5’ non-coding sequence that is otherwise conserved among syngnathids and distantly related percomorphs. In zebrafish, this gene is necessary for outgrowth of the pectoral fin, upstream of *fgf4* and *fgf8* (Nomura et al., 2006), and we show it is expressed similarly in fin margins in a representative percomorph, threespine stickleback (Figure S19) (*Gasterosteus aculeatus*). The eponymous “leaves” of seadragons that are fleshy extensions borne on bony supports extending from the dermal plates, are apparently stiffened by a core of collagenous tissue rather than ossified structures such as fin rays (Figure 1). Homology of the leafy ornaments with fins might pertain only at the level of shared genetic pathways, such as in the case of a cooption or re-deployment of FGF-signaling to elicit proliferation and outgrowth of these superficially fin-like structures. A role in scale or dermal plate patterning for *fgf16* is not known from teleosts, though in birds the orthologous gene, potentially acting with *fgfr1*, can suppress both *shh* expression and downy feather elongation when locally overexpressed (Chen et al., 2016). The seadragon-specific changes in a known AER and integument-patterning gene are intriguing in the context of this morphological oddity, the leafy “appendages”. The seahorse fgf16 protein, we found, bears substitutions in the same amino acid motif deleted in seadragons. The fact that two lineages that have evolved elaborate bony and fleshy ornaments (seadragons and seahorses) both show divergence in a deeply conserved motif of this gene is tantalizing and warrants further comparative work.

## SUMMARY

The genome models of the leafy and weedy seadragon provide a first step towards unlocking how the fantastical forms of these animals and their close relatives have evolved. Even with our preliminary analyses we have identified likely recent expansions of transposable elements, with hypothesized involvement in significant changes to gene families of known consequence, the most surprising being the loss of fgf3 and fgf4 in the sygnathid lineage. Repetitive DNA, including that generated by the activity of TEs, can precipitate genomic rearrangements such as deletions, duplications, and inversions (reviewed in Bourque et al., 2018; Schrader & Schmitz, 2019). It is therefore likely that historical and possibly continuing TE activity in genomes of the syngnathid lineage has resulted in alterations to genome structure and content, with extreme consequences for developmental genetic programs that are otherwise deeply conserved across vertebrates.

## ACKNOWLEDGEMENTS

We are truly grateful for the dedicated efforts of L. Matsushige and the seadragon husbandry staff of the Birch Aquarium at Scripps, who preserved *P. eques* samples crucial for this work. We also thank the Tennessee Aquarium, and particularly Aquarist K. Hurt for generous sharing of *P. taeniolatus* samples. We are indebted to M. Weitzman and D. Turnbull from the UO GC3F for critical assistance with library preparation and sequencing. We would like to acknowledge staff at Carl Zeiss Microscopy, LLC, particularly J. Mancuso, A. Browning, K. Skinner, and R. White for collaborating to scan the Gulf pipefish in Figure S2. We are grateful for T. Desvignes for guidance on the miRNA annotations. We especially thank C. Kimmel for his helpful comments on the manuscript. This work was funded by National Institutes of Health grants RR032670 and P50GM098911 (to WAC), National Science Foundation grant OPP-2015301 (to WAC, SB, and CMS), and Oregon Research Excellence Funds (to WAC). HMH was supported by the Genetics Training Program (NIH T32GM007413) at UO. Purchase of the Zeiss Xradia 620 Versa at the University of Oregon was made possible with funding support from the M.J. Murdock Charitable Trust Grant SR-201812008 (to WAC).

## Data Accessibility

All raw sequencing data associated with this study will be available upon peer-reviewed publication via the NCBI Sequencing Read Archive (SRA), under BioProjects PRJNA765699 and PRJNA765702. Annotation and summary files not in the Supplementary Information will be available upon peer-reviewed publication in the Dryad repository.

