## Supplemental Information for "Leafy and Weedy Seadragon Genomes Connect Genic and Repetitive DNA Features to the Extravagant Biology of Syngnathid Fishes" (from another version of this preprint: version 1, 2021-09-25)

__Supplemental Figures, Tables, and Files__

**SUPPLEMENTAL FIGURES:**

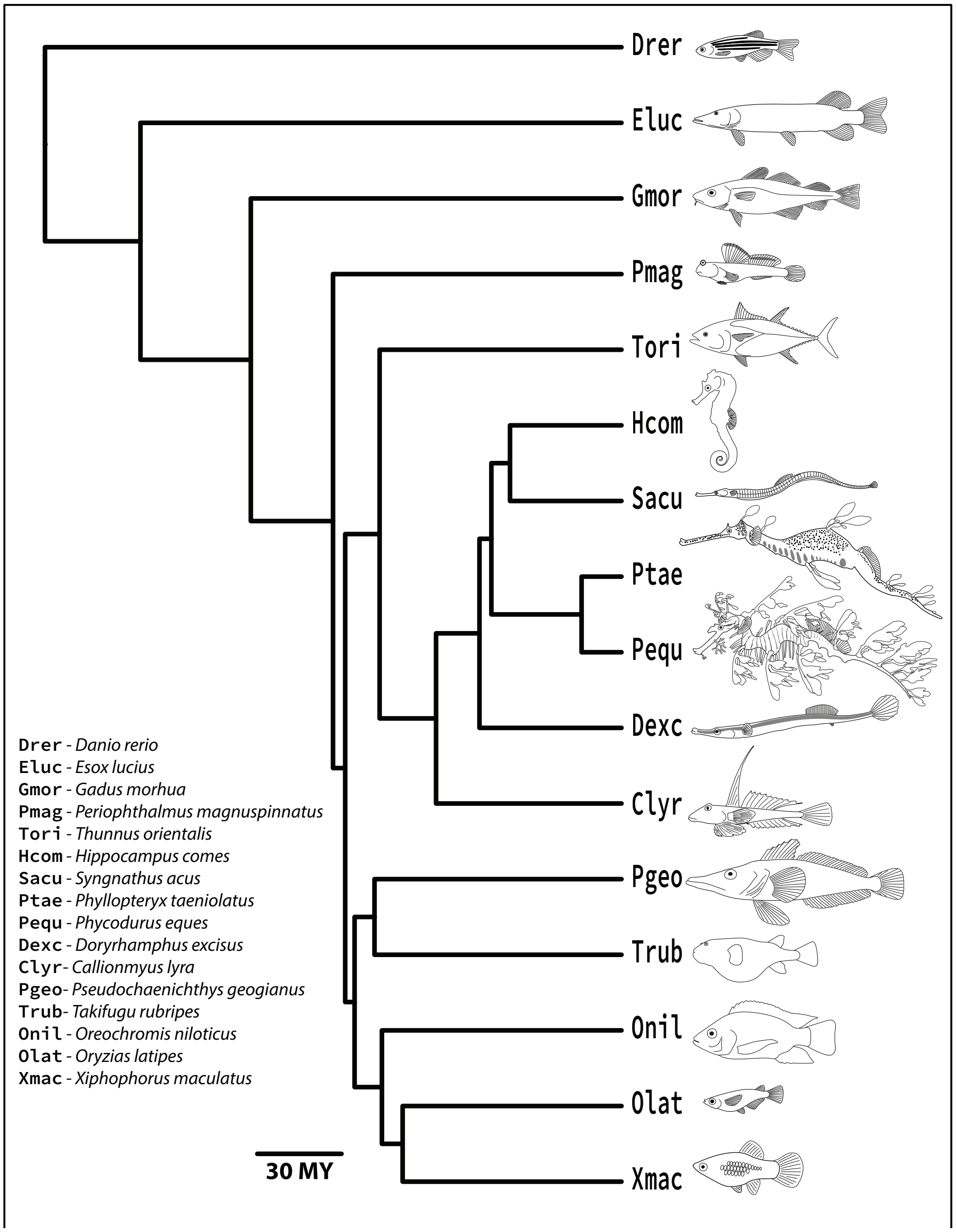


**Figure S1.** Genomes of 16 teleost species provide phylogenetic context for the contributions of repetitive DNA to genome evolution. Represented are evolutionary relationships among a sample of diverse teleosts, according to a time-calibrated phylogenetic tree adapted from Rabosky et al (2018). A

major aim with the selection of these species was to choose genomes assembled with long-read (or at the very least linked-read) technology to the extent possible, to minimize exclusion of repetitive regions from our analysis.

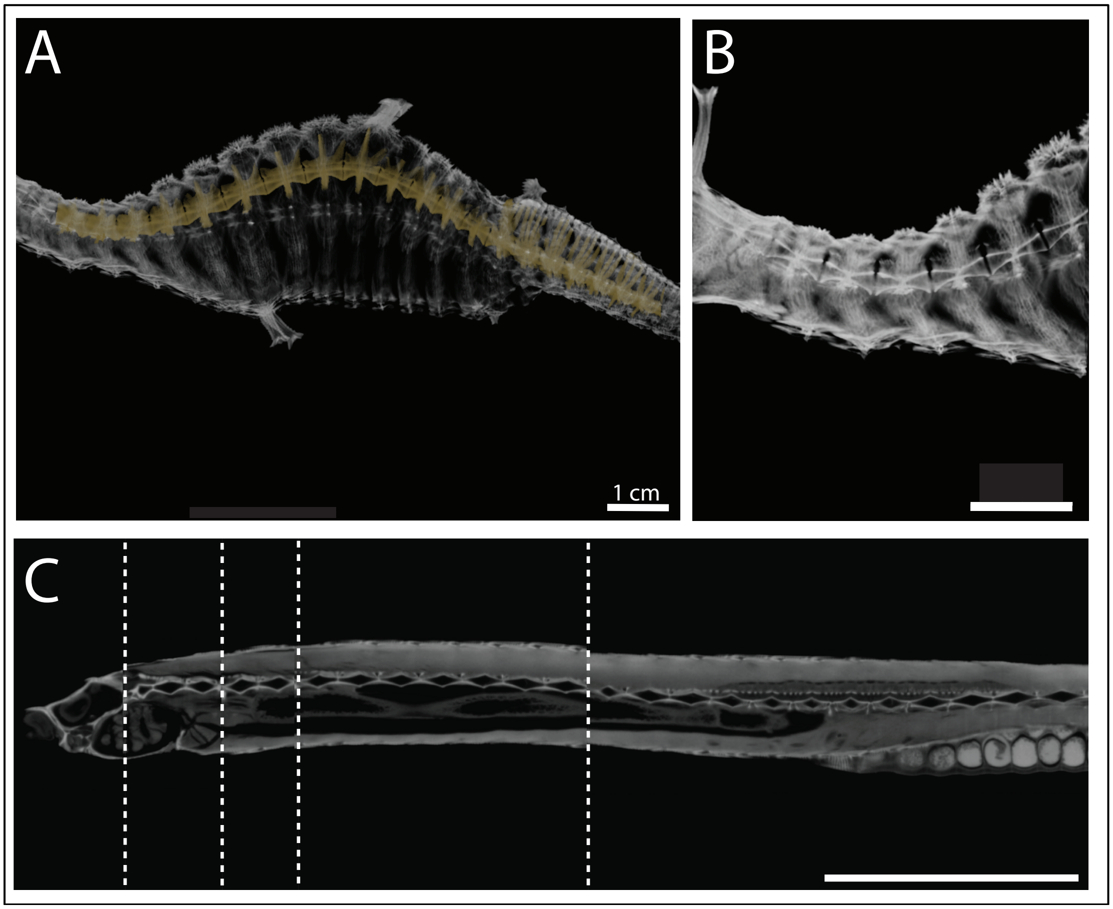


**Figure S2.** Vertebral shape contributes to spinal curvature in weedy seadragon. A) a lateral view of the seadragon with the vertebrae highlighted. B) a lateral detail of the keystone shaped vertebrae close to the head. C) The vertebrae of the straight-bodied Gulf pipefish (Syngnath*us scovelli*) are relatively homogenous in shape. Because of slight lateral curvature of the scanned specimen, a pregnant male, we show here a composite image, with vertical dashed lines marking junctions between the slightly different optical sections needed to follow a midline sagittal plane along the fish. Scale bars = 1 cm.

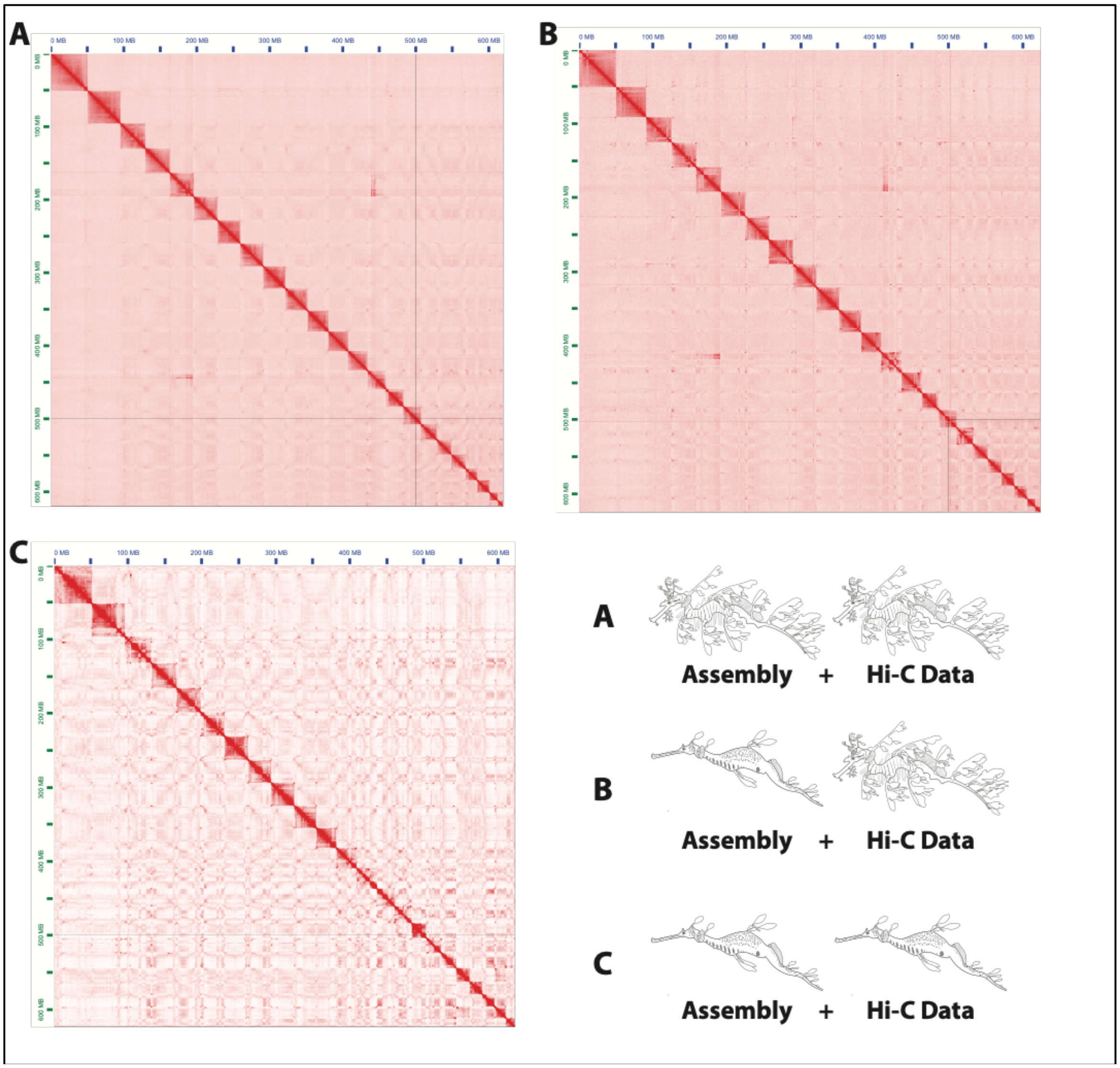


**Figure S3.** Hi-C sequencing of proximity-ligated DNA fragments facilitates the organization of PacBio contigs into seadragon chromosome models. Shown are contact maps, produced by Juicebox (DURAND *et al.* 2016) reflecting the frequency of DNA proximity events across the 23 putative chromosomes after Hi-C scaffolding. Final versions of the leafy and weedy seadragon assemblies (A and B) were each scaffolded using Hi-C data generated from the leafy seadragon. Hi-C data were sequenced for the weedy seadragon, but scaffolding that assembly using the matched species data was problematic (C), possibly due to inferior library preparation.

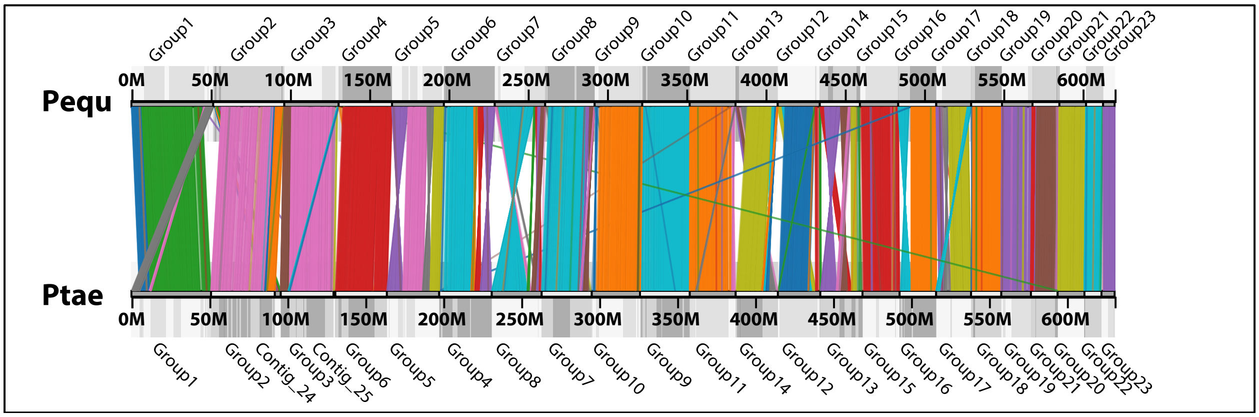


**Figure S4.** The assembled genomes of leafy (Pequ) and weedy (Ptae) seadragons have highly conserved gene synteny on a fine scale. Strings, colored by synteny cluster, connect orthologous genes and groups of genes. A "cluster" is a block of orthologous genes that occur in the same syntenic neighborhood in both genomes, so large blocks of a single color show regions of highly parallel contiguity of orthologs. Hi-C chromosomes (and in two Ptae cases, large Hi-C scaffolds unassigned to chromosomes) are colored in alternating light and dark grey bars, and contig boundaries are indicated by alternating light and dark grey shaded boxes.

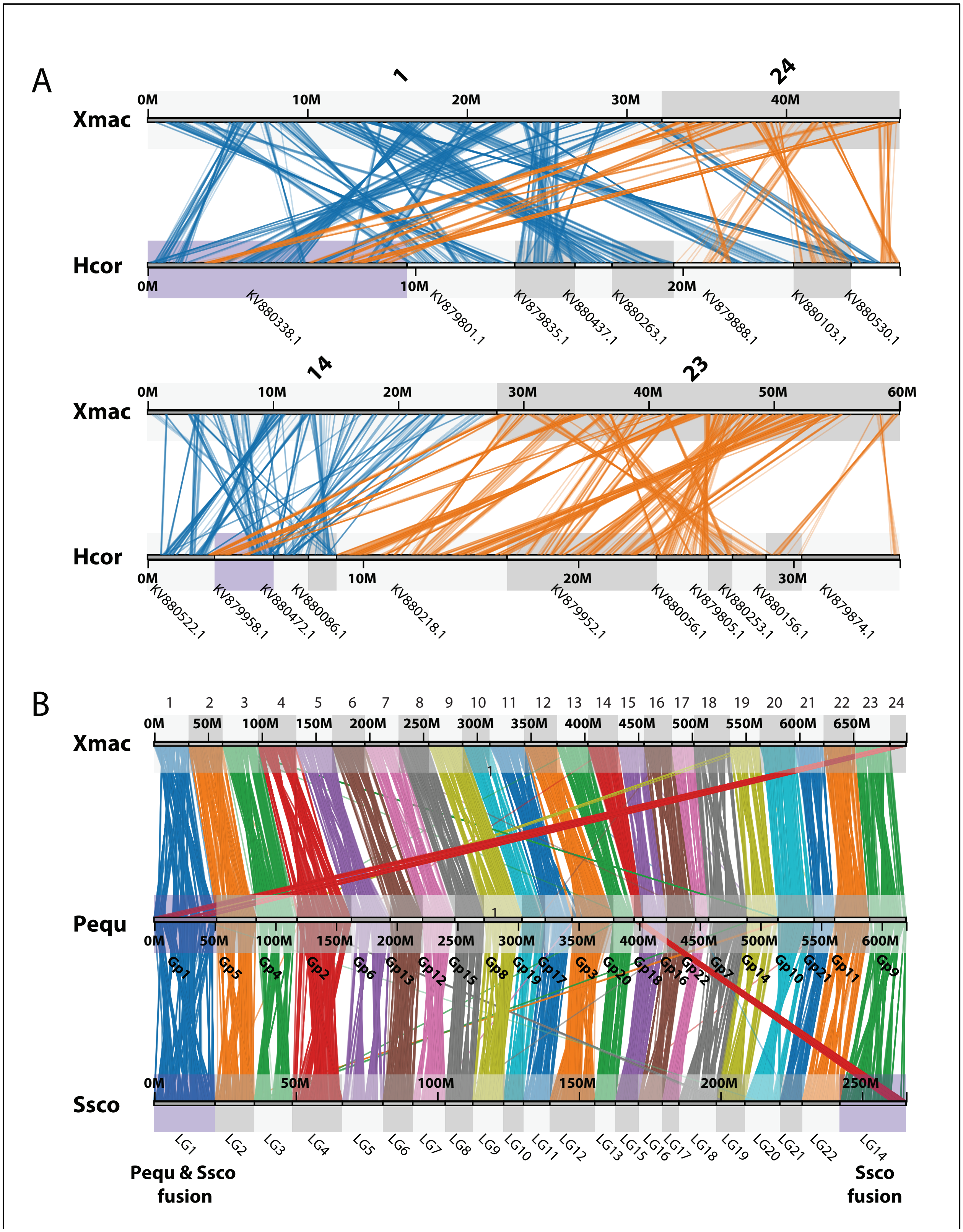


**Figure S5.** The evolutionary sequence of chromosomal fusion events in syngnathid lineages is illustrated by conserved gene synteny comparisons. Fusion of ancestral chromosomes 1 and 24 (numbered as in platyfish) occurred in a common ancestor of seadragons, seahorses, and *Syngnathus* pipefish, but the seahorse+pipefish fusion of chromosomes 14 and 23 (also numbered as in platyfish) occurred after the seadragon lineage diverged. Blocks of conserved gene orthologs are connected by colored strings (A) between the genomes of platyfish (Xmac) and tiger tail seahorse (Hcom), and (B) between platyfish, leafy seadragon (Pequ), and Gulf pipefish (Ssco). Strings are colored by platyfish chromosome. Xmac chromosomes, Pequ Hi-C chromosomes ("Group"), Ssco linkage groups ("LG"), and Hcom scaffolds are demarcated by alternating light and dark shaded bars. A) Two assembled seahorse scaffolds (shaded in lavender) appear to capture the junctions of ancestral chromosomes that became fused in the lineage leading to both seahorses and *Syngnathus* pipefish. B) Leafy seadragon and Gulf pipefish share the fusion of ancestral chromosomes 1 and 24, but the seadragon orthologs of chromosomes 14 and 23 remain separate.

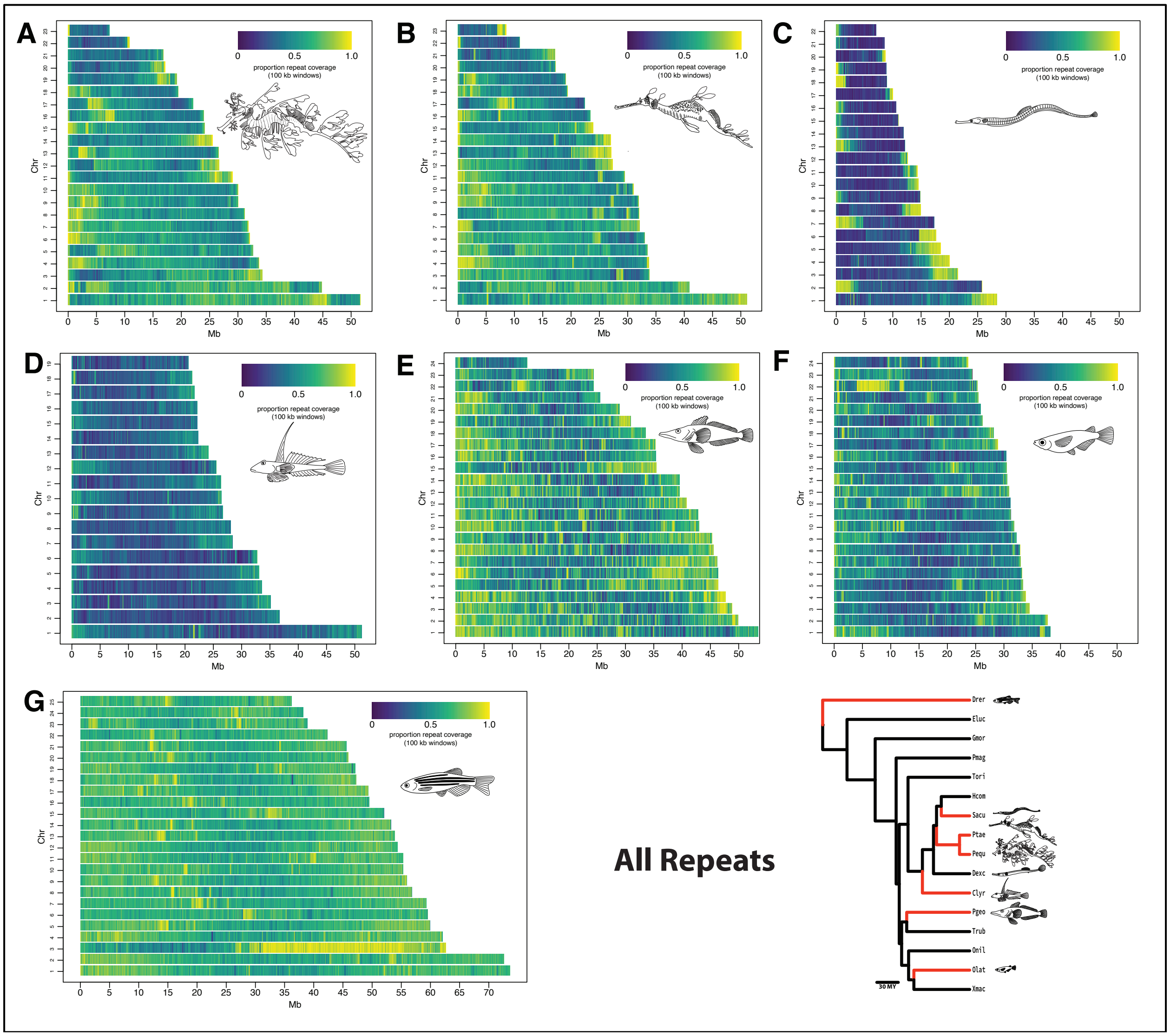


**Figure S6.** Chromosomal variation in repeat density among seven chromosome-scale teleost genome assemblies. Overall repeat basepair occupation of 100 kb windows (expressed as a proportion) is presented as a heatmap for A) leafy seadragon, *Phycodurus eques*; B) weedy seadragon, *Phyllopteryx*

*taeniolatus*; C) greater pipefish, *Syngnathus acus*; D) common dragonet, *Callionymus lyra*; E) South Georiga icefish, *Pseudochaenichthys georgianus*;  F) Japanese medaka, *Oryzias latipes*; and G) zebrafish, *Danio rerio*.

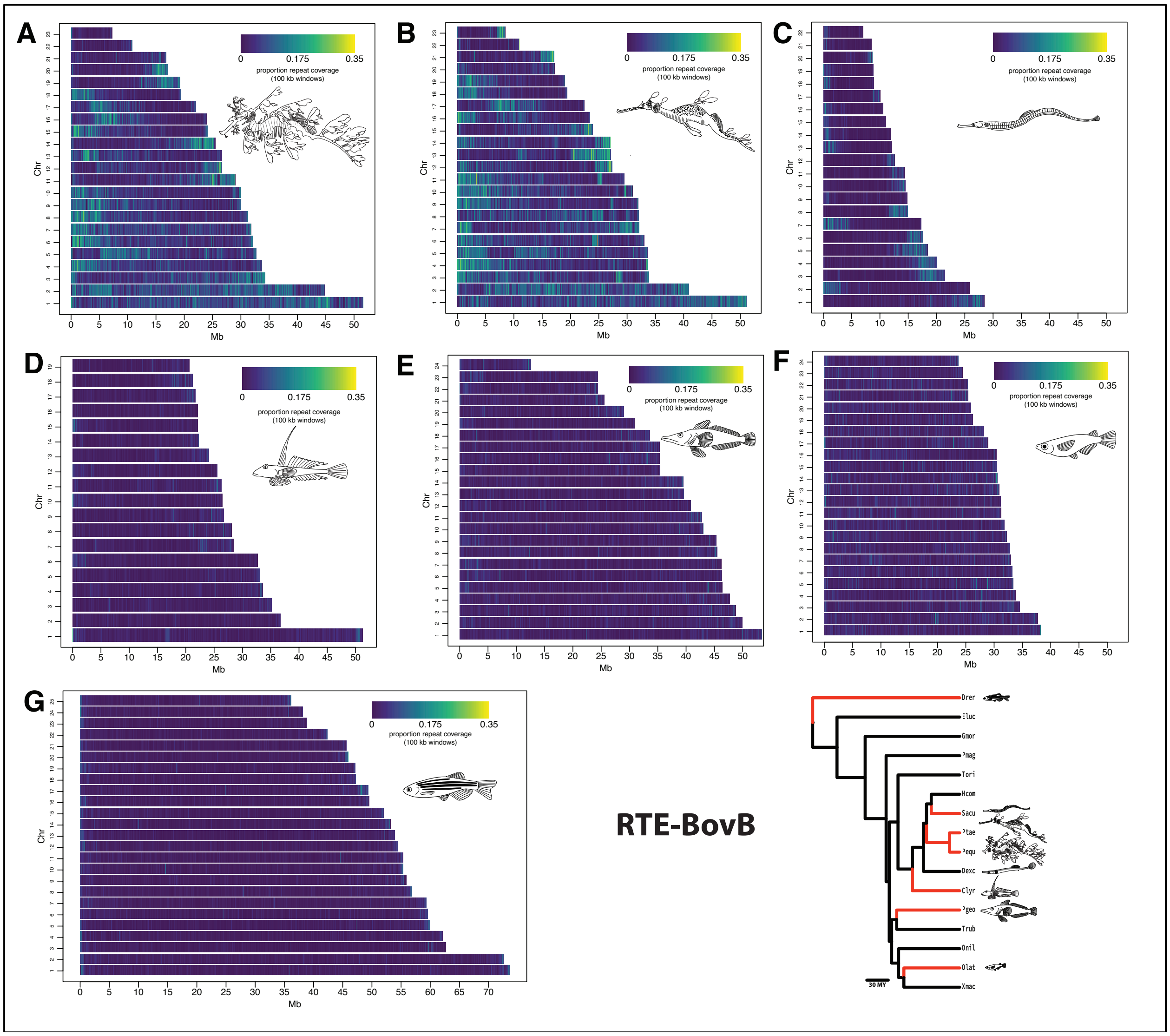


**Figure S7.** Chromosomal variation in the density of RTE-BovB (a LINE retrotransposon) repeats among seven chromosome-scale teleost genome assemblies. BovB basepair occupation of 100 kb windows (expressed as a proportion) is presented as a heatmap for A) leafy seadragon, *Phycodurus eques*; B) weedy seadragon, *Phyllopteryx taeniolatus*; C) greater pipefish, *Syngnathus acus*; D) common dragonet, *Callionymus lyra*; E) South Georgia icefish, *Pseudochaenichthys georgianus*;  F) Japanese medaka, *Oryzias latipes*; and G) zebrafish, *Danio rerio*. BovB repeats are particularly abundant in seadragon genomes.

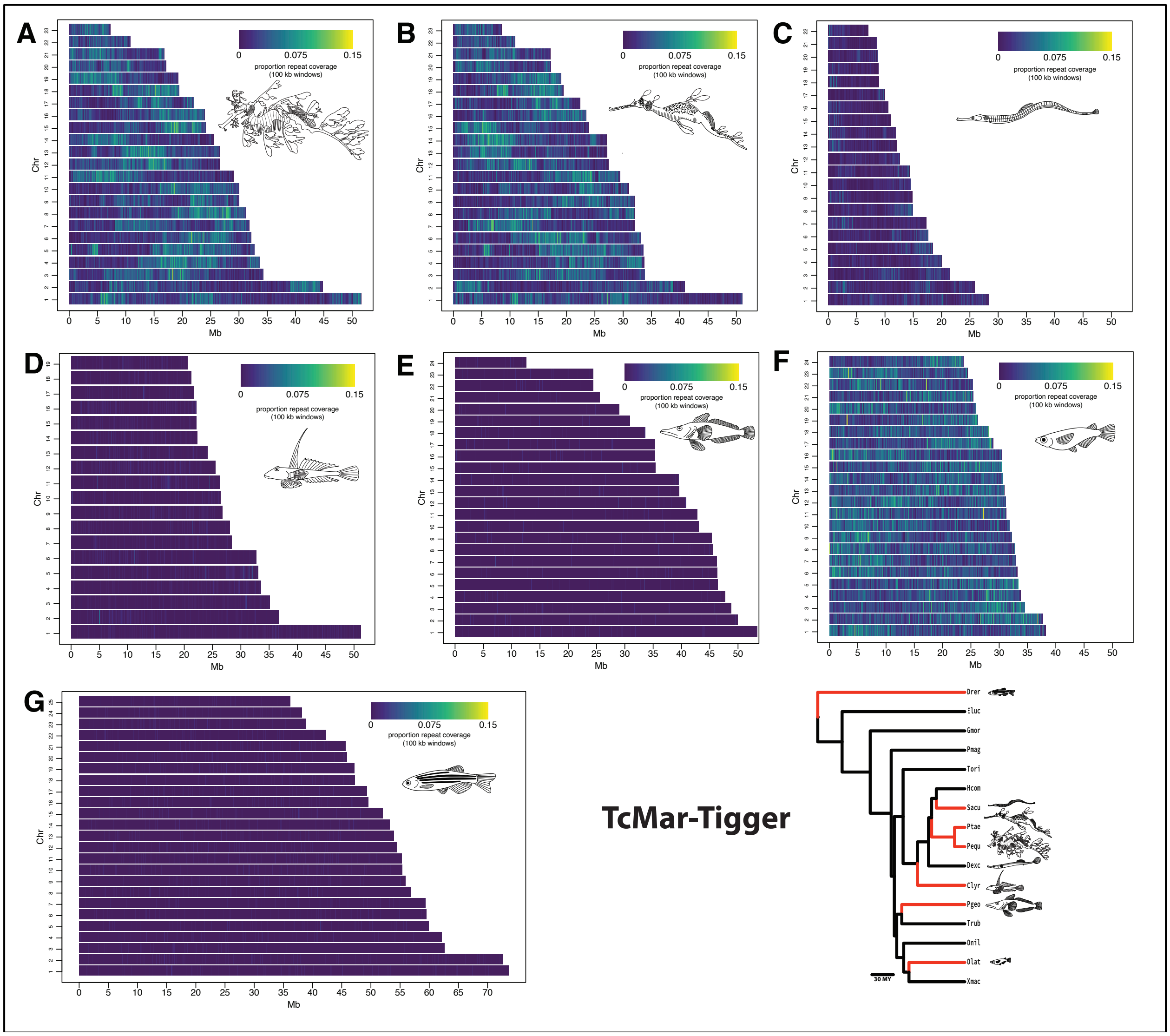


**Figure S8.** Chromosomal variation in the density of TcMar-Tigger (a DNA transposon) repeats among seven chromosome-scale teleost genome assemblies. Tigger basepair occupation of 100 kb windows (expressed as a proportion) is presented as a heatmap for A) leafy seadragon, *Phycodurus eques*; B) weedy seadragon, *Phyllopteryx taeniolatus*; C) greater pipefish, *Syngnathus acus*; D) common dragonet, *Callionymus lyra*; E) South Georiga icefish, *Pseudochaenichthys georgianus*; F) Japanese medaka, *Oryzias latipes*; and G) zebrafish, *Danio rerio*. Tigger repeats are particularly abundant in syngnathid genomes, as well as in other genomes such as medaka. Also note the "patchy" distribution of Tigger repeats in seadragon and greater pipefish genomes, in regions that are largely incongruent with generally high repeat density in those genomes.

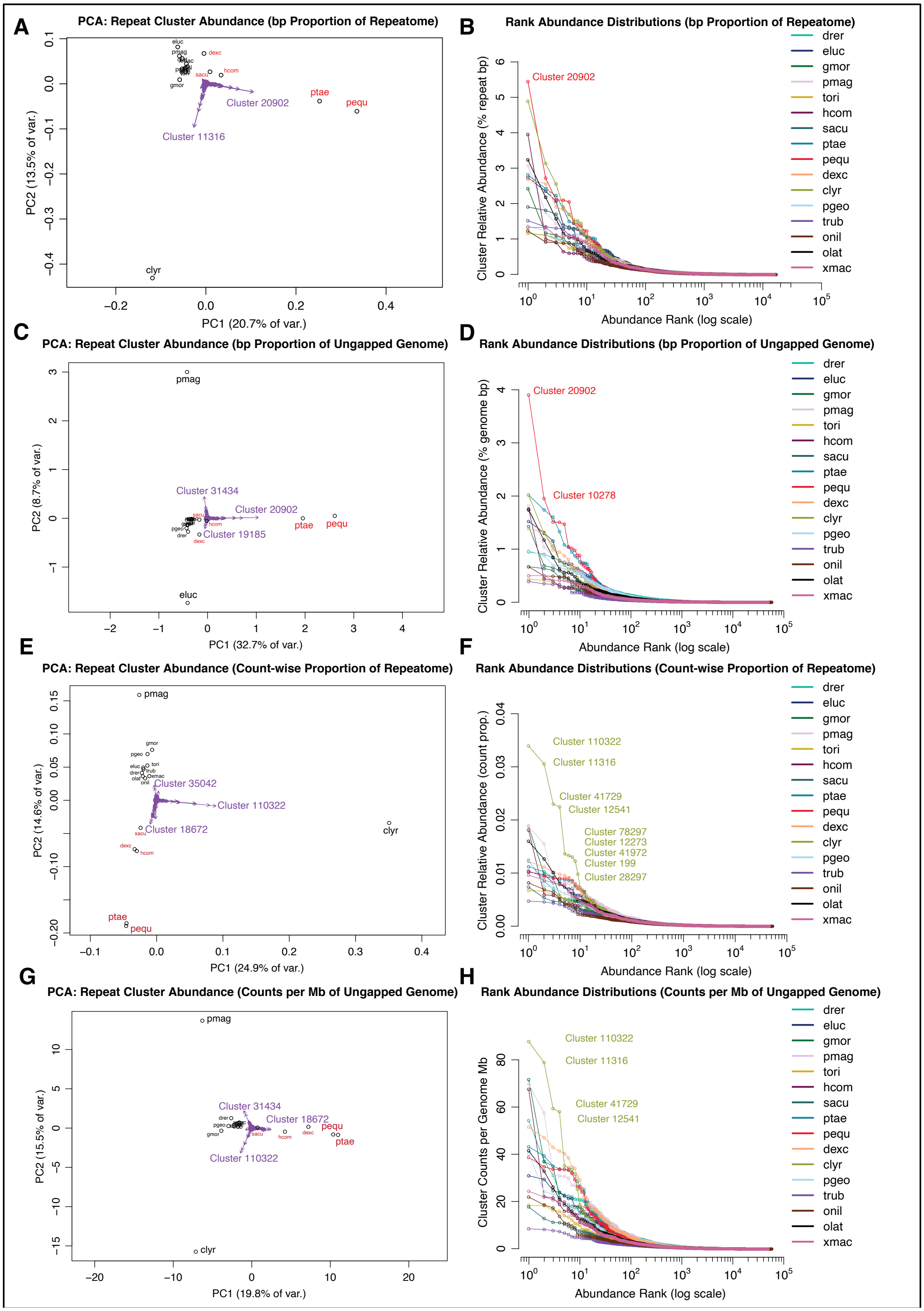


**Figure S9.** Alternative methods for quantifying repeat cluster abundance consistently reveal uniqueness of seadragon repeatomes. Shown are principal components analysis (PCA) bi-plots in the left column of panels, and rank abundance distributions (RADs) in the right column. Repeat clusters were quantified as

(A-B) the relative proportion of total repeat basepairs, (C-D) the relative proportion of the entire genome basepairs, (E-F) the count-wise proportion of total repeat counts, and (G-H) as repeat counts per Mb of genome. In each plot pair, individual repeat clusters that load heavily on PCs or are especially abundant in individual genomes are noted in purple (PCA plots) or the matching species color (RAD plots).

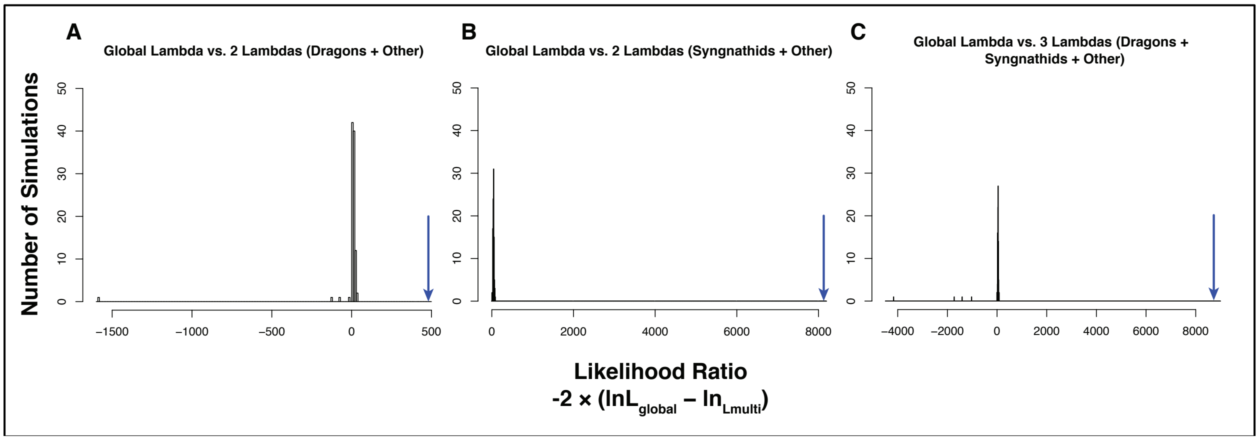


**Figure S10.** Evolutionary simulations with CAFE 5 (Mendes *et al.* 2020) provide evidence for seadragon and syngnathid branch-specific gene family expansion/contraction rates ($\lambda$) that are distinct from a global estimate for the 21-species tree. We compared the likelihood for a single-$\lambda$ model to that of a multi-$\lambda$ model in which branches of interest had a separate, distinct $\lambda$, using the likelihood ratio (-2 × ($lnL_{global} - lnL_{multi}$)). For each test we compared the likelihood ratio observed from the data to a distribution based on 100 Brownian Motion gene family evolution simulations. Specific branch tests included (A) a distinct $\lambda$ for the internal seadragon ancestor branch, (B) a distinct $\lambda$ for the internal syngnathid ancestor branch, and (C) distinct $\lambda$s for both the seadragon and syngnathid ancestor branches. Shown are histograms (gray bars) of the simulated likelihood ratios, and blue arrows marking the LRs observed from the actual data.

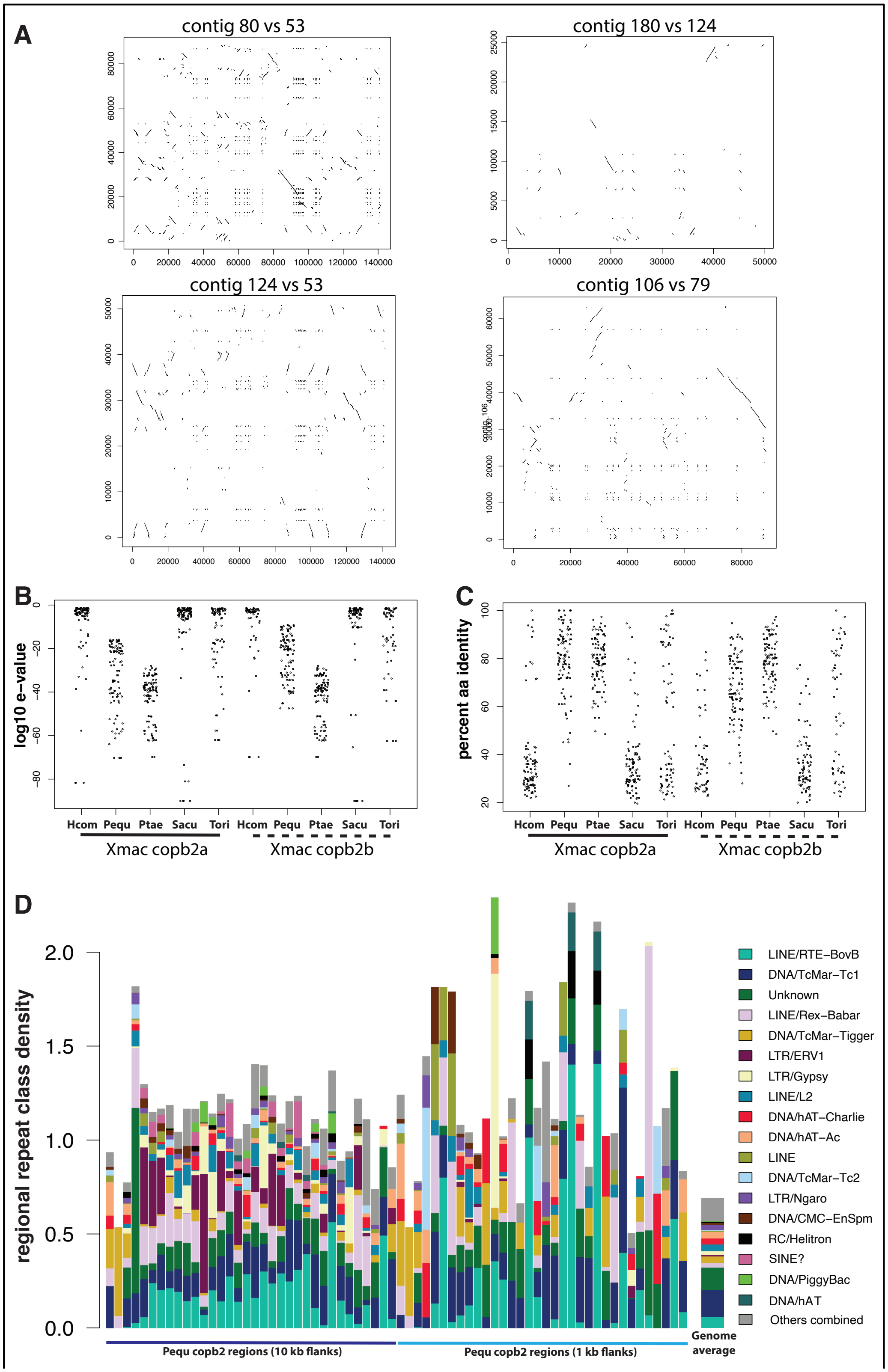


**Figure S11.** Leafy and weedy seadragon genome assemblies provide evidence for expansions of *coatomer subunit beta 2* (*copb2*) genes. A) Dotplot examples reflecting genomic sequence alignments of regions from the weedy seadragon genome containing putative *copb2* copies. The dotplots do not show large tracks of conserved sequence between regions, suggesting that they are not alternative haplotypes assembled separately. B) Distributions of e-values for tblastn queries of platyfish (Xmac) *copb2* paralogs against tiger tail seahorse (Hcom), leafy and weedy seadragons (Pequ and Ptae), greater pipefish (Sacu), and Pacific bluefin tuna (Tori) genomes. C) Distributions of amino acid percent identity for tblastn queries of *X. maculatus* (platyfish) *copb2* paralogs against the genomes listed in C. Each point in B and C represents an individual blast high-scoring pair (hsps). D) Repeat landscapes in *copb2* regions and flanking sequence within the leafy seadragon genome. Each region reflects an individual gene (or pseudogene), and either 10 kb flanking sequence in both directions (first 34 bars) or 1 kb flanking sequence in both directions (second 34 bars). Colors in stacked bars reflect summed basepairs annotated for a specific repeat class divided by the total length of the region. The last, thick bar reflects mean repeat class abundances from 499 random samples of 34 genes (and their 10 kb flanking regions), used as a genomic “background” for comparison. The background repeat frequencies for 1 kb flanking regions were nearly identical, and are not shown. Bars can exceed y-axis values of 1.0 due to overlapping repeat annotations (e.g., those on opposite strands).

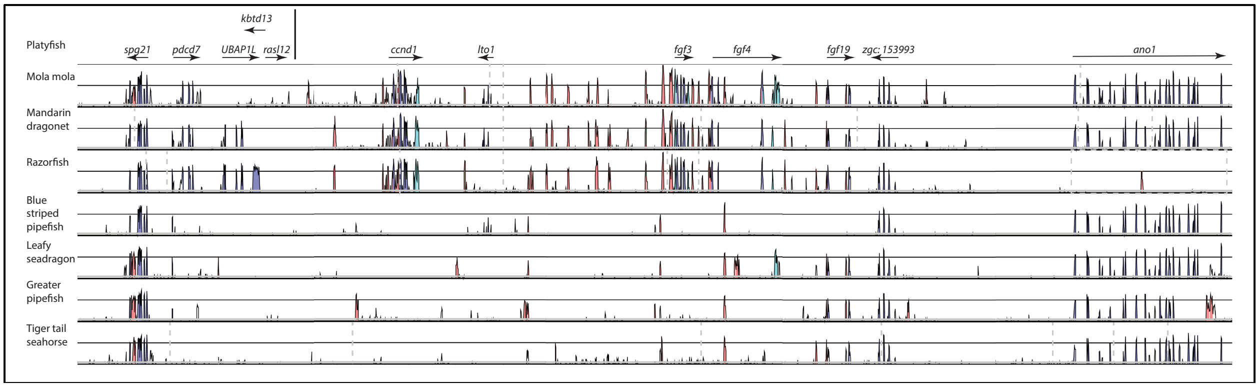


**Figure S12.** Syngnathids have uniquely lost *fgf3, fgf4,* and other neighboring genes. These losses are demonstrated on the vista plot with platyfish as the reference. In the plot, conserved exons are marked in dark blue peaks, conserved non-coding sequences in pink, and conserved three prime untranslated regions in light blue. Gaps in the tiger tail seahorse, mola mola, mandarin dragonet, and razorfish assemblies are noted with grey dashed lines (small gaps) and grey dashed boxes (larger gaps). There is a 150Kb gene sparse region that is removed from the vista and noted with the black bar. Additionally, *ano1* is present on a different contig for razorfish than *fgf3* and *fgf4* due to a fragmented assembly.

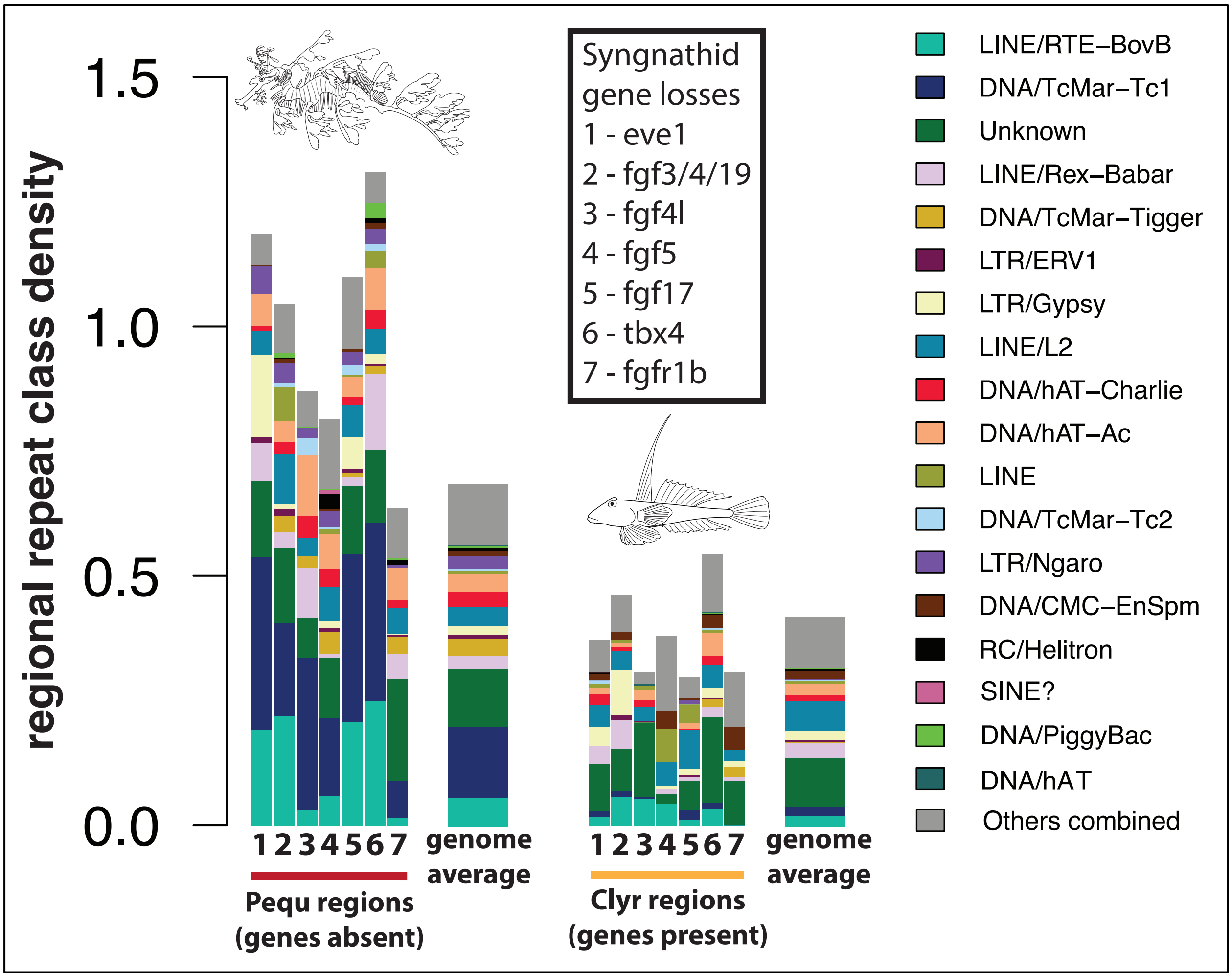


**Figure S13.** Repeat density is elevated in key regions of syngnathid-specific developmental gene loss. Repeat landscapes at seven regions in which highly conserved developmental genes have been lost in syngnathid fishes are shown using repeat density bar plots for leafy seadragon (as a syngnathid representative), and orthologous regions in a close outgroup, the common dragonet. Colors in stacked bars reflect summed basepairs annotated for a specific repeat class divided by the total length of the region. Thick bars reflect mean repeat class abundances from 499 random samples of seven genes (with regions extending to immediate flanking genes in both directions), used as genomic “backgrounds” for comparison. Bars can exceed y-axis values of 1.0 due to overlapping repeat annotations (e.g., those on opposite strands). Overall repeat density is significantly higher in the focal regions relative to the background distribution in the seadragon, but not the dragonet, genome.

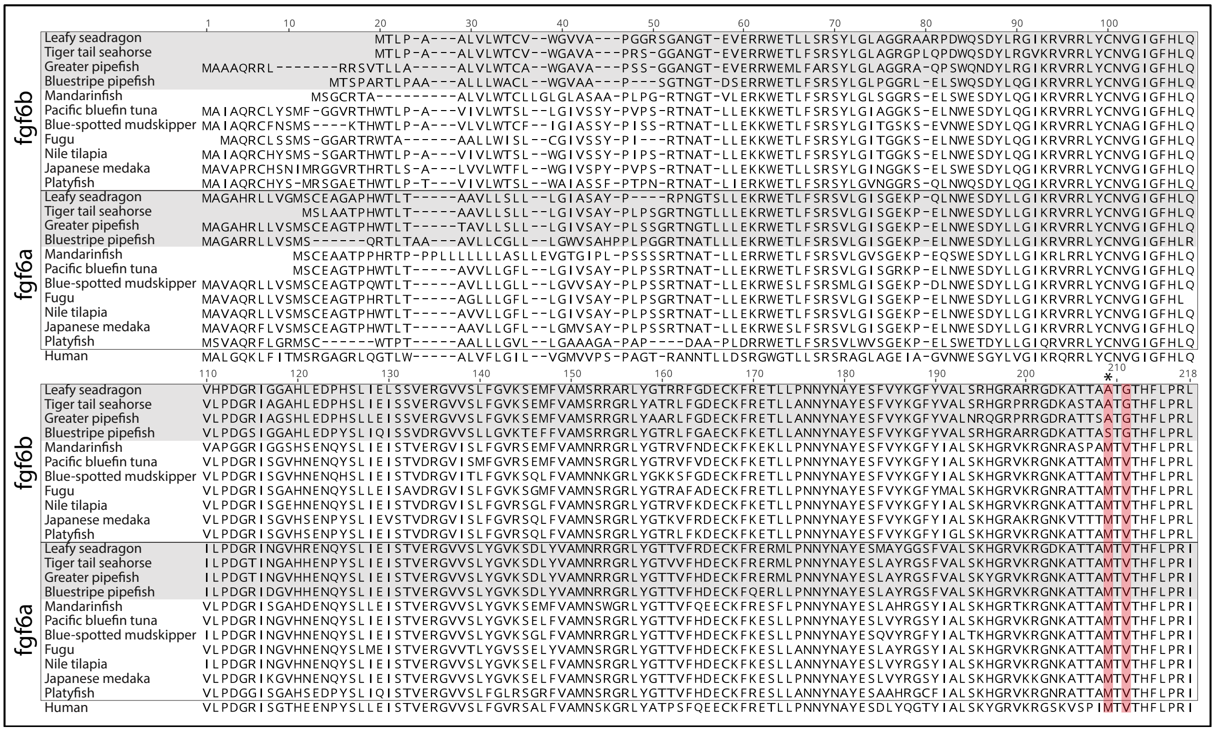

**Figure S14.** *fgf6b* was inferred to have experienced positive selection (before adjusting for false discovery rate) in syngnathids, and one of the derived substitutions in this category is predicted to be deleterious. Aligned Fgf6b protein sequences from percomorph fishes and human are shown, with syngnathid sequences highlighted in grey. Two syngnathid-specific mutations at deeply conserved sites are highlighted in red; the site with the asterisk was predicted to be deleterious by Provean (CHOI AND CHAN 2015). By contrast syngnathid Fgf6a has maintained these conserved amino acids.

| | |
|---|---|
| | * |
| Ola4 | ADFGLARGVHQIDYYKKTTNGRLPVK |
| Tru4 | ADFGLARGVHQIDYYKKTTNGRLPVK |
| Oni4 | ADFGLARGVHQIDYYKKTTNGRLPVK |
| Xma4 | ADFGLARGVHQIDYYKKTTNGRLPVK |
| Peq4 | ADFGLARGVHQIDYYKKTTNGRLPVK |
| Dex4 | ADFGLARGVHQIDYYKKTTNGRLPVK |
| Hco4 | ADFGLARGVHQIDYYKKTTNGRLPVK |
| Xma3 | ADFGLARDVHNIDYYKKTTNGRLPVK |
| Ola3 | ADFGLARDVHNIDYYKKTTNGRLPVK |
| Oni3 | ADFGLARDVHNIDYYKKTTNGRLPVK |
| Tru3 | ADFGLARDVHNIDYYKKTTNGRLPVK |
| Dex3 | ADFGLARDVHNIDYYKKTTNGRLPVK |
| Peq3 | ADFGLARDVHNIDYYKKTTNGRLPVK |
| Oni2 | ADFGLARDVHNIDYYKKTTNGRLPVK |
| Ola2 | ADFGLARDVHNIDYYKKTTNGRLPVK |
| Xma2 | ADFGLARDVHNIDYYKKTTNGRLPVK |
| Tru2 | ADFGLARDVHNIDYYKKTTNGRLPVK |
| Dex2 | ADFGLARDVHNIDYYKKTTNGRLPVK |
| Peq2 | ADFGLARDVHNIDYYKKTTNGRLPVK |
| Hco2 | ADFGLARDVHNIDYYKKTTNGRLPVK |
| Hco3 | ADFGLARDVHNLDYYKKTTNGRLPVK |
| TruA | ADFGLARDIHHIDYYKKTTNGRLPVK |
| OlaA | ADFGLARDIHHIDYYKKTTNGRLPVK |
| XmaA | ADFGLARDIHHIDYYKKTTNGRLPVK |
| OauA | ADFGLARDIHHIDYYKKTTNGRLPVK |
| DexA | ADFGLARDIHHIDYYKKTTNGRLPVK |
| HcoA | ADFGLARDIHHIDYYKKTTNGRLPVK |
| PeqA | ADFGLARDIHHIDYYKKTTNGRLPVK |
| OniB | ADFGLARDIHHIDYYKKTTNGRLPVK |
| OlaB | ADFGLARDIHHIDYYKKTTNGRLPVK |
| TruB | ADFGLARDVHHIDYYKKTTNGRLPVK |
| XmaB | ADFGLARDIHHIDYYKKTTNGRLPVK |
| DexB | ADFGLARDVHHVDYYKKTTNGPLPVK |
| HcoB | ADFGLARDVHHLDYYKKTTNGPLPVK |
| PeqB | ADFGLARDVHHVDYYKKTTNGPLPVK |
| Hsa | ADFGLARDIHHIDYYKKTTNGRLPVK |

**Figure S15.** A unique substitution in the deeply conserved activation loop of syngnathid *fgfr1b*. The activation loop from teleost fish *fgfr* sequences are aligned and shown. The receptor number is appended to the three-letter species code. For *fgfr1* sequences, A and B are appended to the end. The human (Hsa) activation loop sequence for *fgfr1* is included on the bottom row. The syngnathid *fgfr1b* sequences are highlighted in purple, and the syngnathid *fgfr1b* R to P substitution is noted with an asterisk.

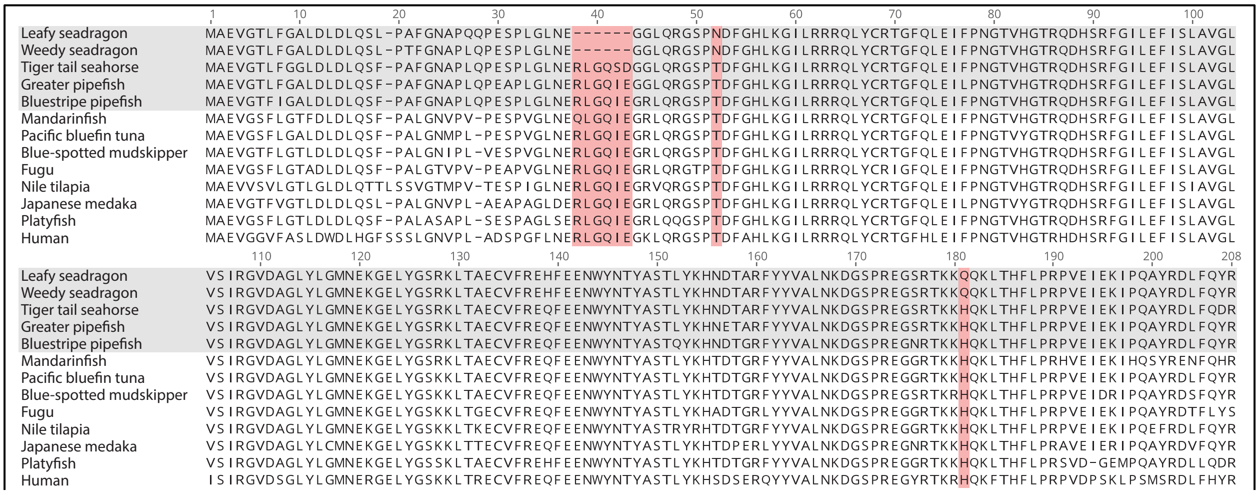


**Figure S16.** An alignment of predicted Fgf16 protein sequences reveals a deletion of six deeply conserved amino acids and substitutions of two other conserved amino acids (shaded in red) in the seadragons. These differences are not shared by the other syngnathids (clade shaded in grey), though seahorse has two substitutions in the 6 amino acid motif.

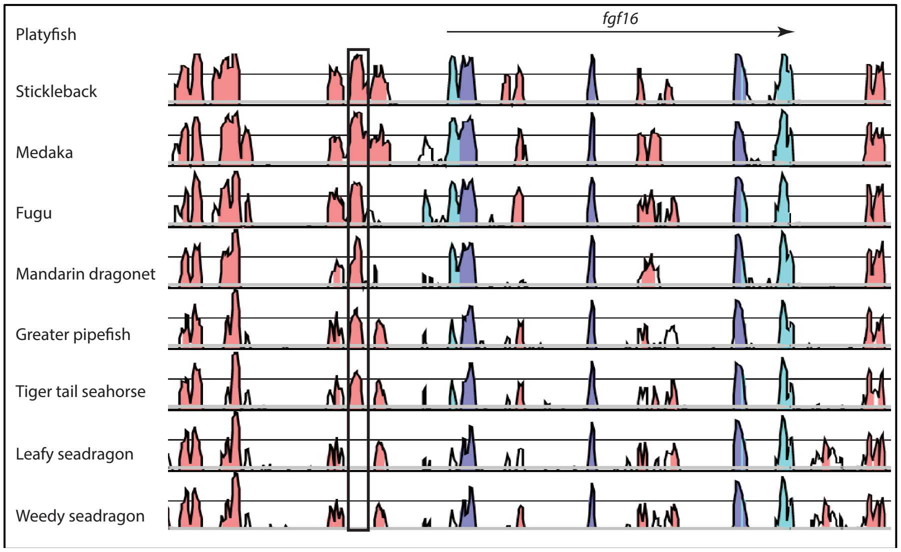


**Figure S17.** Leafy and weedy seadragons have lost a conserved non-coding element (shown in the black box) near *fgf16* that is retained in tiger tail seahorse, greater pipefish, and other percomorph fish. The vista plot highlights conserved non-coding sequences in pink, conserved UTRs in light blue, and conserved exons in dark blue.

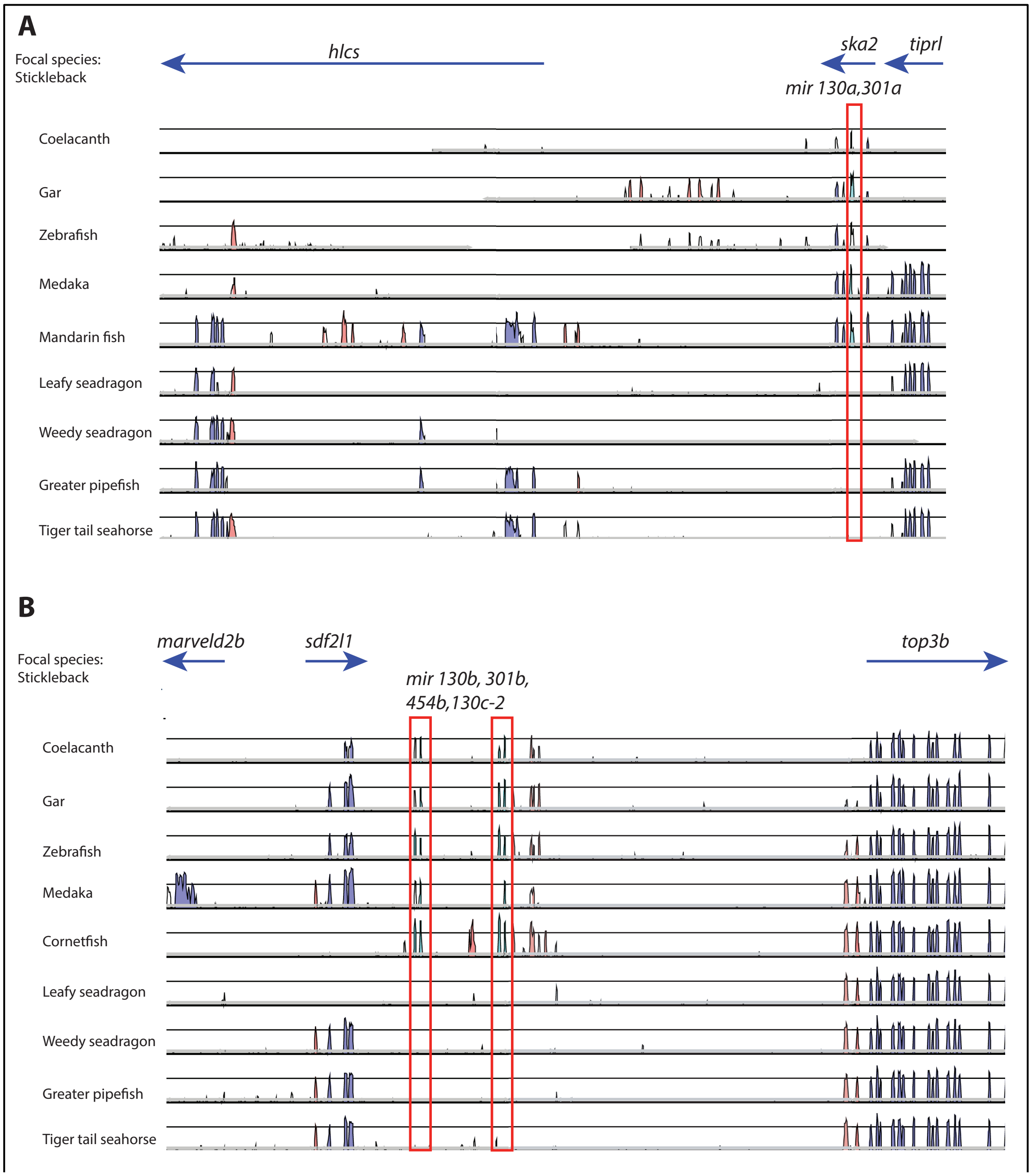


**Figure S18.** Two conserved miRNA clusters appear to be lost in the syngnathid lineage, clusters containing A) *mir130a (*also including *mir301a)* and B) *mir130b* (additionally having *mir301b*, *mir130c-2*, and *mir454b*). Vista plots using threespine stickleback as the reference illustrate these losses (boxed in red). In these plots, conserved exons are demonstrated in dark blue peaks, conserved non-coding elements are noted in pink peaks, and conserved miRNAs are shown in light blue peaks. The orientation (blue arrows) and names of protein coding genes are shown above the plot. Syngnathids have further lost *ska2*, the protein coding gene in whose intron *mir130a* and *mir301a* normally reside (A).

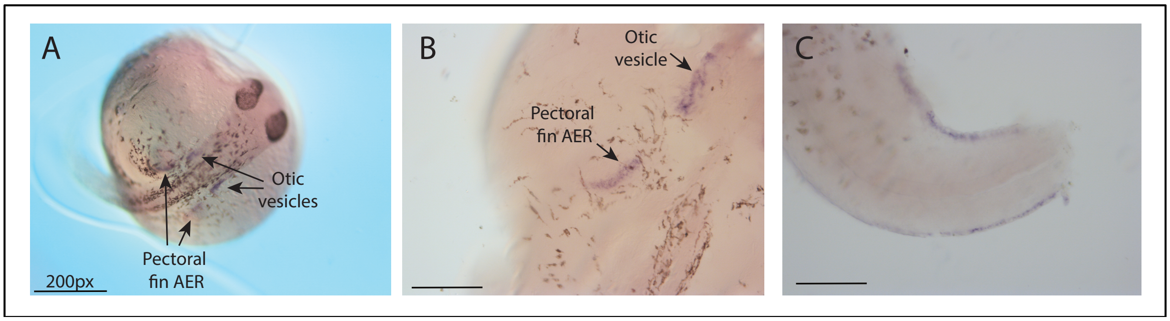


**Figure S19.** We confirmed that *fgf16* is expressed in a percomorph fish in the fin bud AER, as had been described in a phylogenetically distant taxon, zebrafish. Shown is the expression in developing threespine stickleback (*Gasterosteus aculeatus*; 75 hours post fertilization). Panels A) and B) show the expression domains in the marginal tissue of the fin bud and in the developing ear. C) The caudal fin margin also expresses the gene. The anti-sense RNA probe was transcribed from a cloned fragment amplified from stickleback genomic DNA using these primer sequences: forward primer 5' CGGACGAACTGGTTACTACATC3' and reverse 5' TTCATCCCTCGGACAGTCTTC3'.

### SUPPLEMENTAL TABLES:

| | fgf3 | fgf7 | fgf10a | fgf10b | fgf22 | fgf4 | fgf4-like | fgf5 | fgf6a | fgf6b | fgf8a | fgf8b | fgf17 | fgf24 | fgf16 | fgf20a | fgf20b | fgf19 |
|---|---|---|---|---|---|---|---|---|---|---|---|---|---|---|---|---|---|---|
| Leafy Seadragon | - | + | + | + | + | - | - | + | + | + | + | + | - | + | + | + | + | + |
| Tiger tail Seahorse | - | + | + | + | + | - | + | + | + | + | + | + | + | + | + | + | + | + |
| Greater Pipefish | - | + | + | + | + | - | - | - | + | + | + | + | + | + | + | + | + | + |
| Bluestripe Pipefish | - | + | + | + | + | - | + | + | + | + | + | + | - | + | + | + | + | - |

**Supplemental Table 1.** Several developmentally important, conserved FGF ligands have been lost in syngnathid genomes. Shown in this table are the complements for these genes in leafy seadragon, tiger tail seahorse, greater pipefish, and bluestripe pipefish. All share losses of *fgf3* and *fgf4, fgf4-like* and *fgf17* has been lost in the seadragons and greater pipefish, and *fgf5* and *fgf19* are missing in the greater pipefish. Alternating column shading denotes gene subfamily groupings (ITOH AND KONISHI 2007; ORNITZ AND ITOH 2015).

| *Gene* | *Branch.Tested* | *Likelihood.Ratio.Test* | *P.value* | *Adjusted.P.value* | *Sites.Detected.Under.Positive.Selection* | *adjustP* |
|---|---|---|---|---|---|---|
| fgfrl1a | syngnathid | 1.70 | 0.19 | 1.00 | | 1.00 |
| fgfrl1b | syngnathid | 0.00 | 1.00 | 1.00 | | 1.00 |
| fgf8a | syngnathid | 0.00 | 1.00 | 1.00 | | 1.00 |
| fgf8b | syngnathid | 0.00 | 1.00 | 1.00 | | 1.00 |
| fgf10a | syngnathid | 0.00 | 1.00 | 1.00 | | 1.00 |
| fgf10b | syngnathid | 0.03 | 0.86 | 1.00 | | 1.00 |
| fgf19 | syngnathid | 0.00 | 1.00 | 1.00 | | 1.00 |
| fgfr1a | syngnathid | 0.00 | 1.00 | 1.00 | | 1.00 |
| fgfr1b | syngnathid | 0.07 | 0.79 | 1.00 | | 1.00 |
| fgfr2 | syngnathid | 0.00 | 0.97 | 1.00 | | 1.00 |
| fgfr3 | syngnathid | 0.00 | 1.00 | 1.00 | | 1.00 |
| fgfr4 | syngnathid | 0.00 | 1.00 | 1.00 | | 1.00 |
| fgf17 | syngnathid | 11.12 | 0.00 | 0.02 | 18, 19, 20, 60, 120 | 0.03 |
| fgf6b | syngnathid | 4.70 | 0.03 | 0.79 | 25,161,170 | 0.85 |
| fgf6a | syngnathid | 0.00 | 1.00 | 1.00 | | 1.00 |
| fgf20a | syngnathid | 0.00 | 1.00 | 1.00 | | 1.00 |
| fgf20b | syngnathid | 0.00 | 1.00 | 1.00 | | 1.00 |
| fgf4-like | syngnathid | 0.00 | 1.00 | 1.00 | | 1.00 |
| fgf5 | syngnathid | 0.00 | 1.00 | 1.00 | | 1.00 |
| fgf22 | syngnathid | 0.00 | 1.00 | 1.00 | | 1.00 |
| fgf7 | syngnathid | 0.00 | 1.00 | 1.00 | | 1.00 |
| fgf24 | syngnathid | 0.00 | 1.00 | 1.00 | | 1.00 |
| fgfr1a | Peq/Hco | 0.00 | 1.00 | 1.00 | | 1.00 |
| fgfr1b | Peq/Hco | 5.72 | 0.02 | 0.45 | 3,35,47,52,56,90,97,125,127,128,129,137,170,293,355,406,407 | 0.49 |
| fgf20a | Peq/Hco | 0.00 | 1.00 | 1.00 | | 1.00 |
| fgf20b | Peq/Hco | 0.00 | 1.00 | 1.00 | | 1.00 |
| fgf16 | Peq | 0.00 | 1.00 | 1.00 | | 1.00 |
| fgf6b | Peq | 0.00 | 1.00 | 1.00 | | 1.00 |
| fgf6a | Peq | 0.00 | 1.00 | 1.00 | | 1.00 |
| fgf16 | Hco | 0.00 | 1.00 | 1.00 | | 1.00 |

**Supplemental Table 2.** Subtle evidence for positive selection on FGF ligands and receptors in syngnathid fishes is revealed by phylogenetic analysis by maximum likelihood (PAML). Included are PAML branch site model results for each gene and the foreground branches tested. *fgf17* and *fgf6b* show potential signatures of positive selection in the lineage leading to syngnathids, and *fgfr1b* independently in lineages leading to leafy seadragon and tiger tail seahorse, although two of these fall within the range of false discovery after multiple testing is considered.

**SUPPLEMENTAL FILE CAPTIONS (See individual spreadsheet files):**

**Supplemental File 1.** The 20 most abundant repeat clusters in leafy and weedy seadragon genomes, as measured by basepair proportion of ungapped genome length. Included are the cluster IDs, % abundance values, abundance ranks, consensus class, and total bp in seadragon genomes (both species) and the remaining taxa, for each repeat cluster. Clusters highlighted in orange and green loaded heavily on major PCA axes differentiating seadragons from other teleosts, based on basepair- and count-wise abundance measures, respectively.

**Supplemental File 2.** Statistically significant expansions/contractions of gene families from the CAFE 5 analysis. Sheet A) Branches of interest, including all terminal syngnathiform branches, the seadragon ancestral branch, and the syngnathid ancestral branch, are included here. Gene families statistically significantly expanded/contracted along the terminal seadragon branches; the ancestral seadragon branch, and the syngnathid ancestral branch are highlighted in red and blue, respectively. Numbers in cells indicate the estimated numbers of family members gained or lost (-) along each branch. Sheet B) Results from a KEGG pathway overrepresentation test for gene families expanded/contracted along the leafy seadragon branch, with the FDR controlled at 0.1. Pathways highlighted in green do not overlap with those significantly overrepresented in the list of "background" rapidly evolving gene families. Sheet C) KEGG pathway overrepresentation test results for expanded/contracted families in the weedy seadragon lineage. Sheet D) KEGG pathway overrepresentation test results for expanded/contracted families along the branch leading to seadragons. Sheet E) KEGG pathway overrepresentation test results for expanded/contracted families along the branch leading to syngnathids.

**Supplemental File 3.** Putative *copb2* duplicate regions in leafy and weedy seadragon genomes based on tblastn searches using 2 *X. maculatus* (paralog) queries. Sheet A) Regions of the leafy seadragon genome with one or more high-scoring pairs from the blast search. Sheet B) Similar information as in Sheet A, except that overlapping hits resulting from the two *X. maculatus* paralogs have been merged to eliminate redundancy, and coordinates have been adjusted to include 1 kb of flanking sequence on each side when possible. Sheets C-D) Information as in Sheets A-B, but for weedy seadragon. One of the *copb2* paralogs (on *P. eques* scaffold 13 and *P. taeniolatus* scaffold 14) is likely orthologous to the *X. maculatus copb2* paralog on chromosome (Xmac 6), and the others are possibly derived from this sequence. Echoing one of our other findings - that the fgf3/4/19 locus could harbor some inherent structural instability in the syngnathid lineage - leafy seadragon sequences for *copb2* fall in three locations on scaffold 2, in a region that also harbors a set of genes (e.g., *selenos*, *chsy1*, *snrpa1*) whose orthologs are adjacent to the *fgf3/4/19* cluster region in outgroup percomorphs (e.g., platyfish and threespine stickleback), but in syngnathids fall many megabases away from the remains of the cluster.

**Supplemental File 4.** Statistical enrichment test results for elevated repeat density in *cop2b* regions (leafy seadragon genome) and in seven syngnathid-specific gene loss regions (leafy seadragon and common dragonet genomes). Columns include the repeat class tested, the observed and randomly resampled among-region means, 95% confidence intervals for the resampled means, the ratio of observed to randomly resampled means ("density fold change"), and test p-values (original and FDR-adjusted). Clusters significantly enriched (FDR=0.1) in focal regions relative to randomly sampled regions are highlighted in green. Repeat classes with observed mean regional densities > 10% are in red font. Sheet A) Enrichment test results for *copb2* regions with 10 kb flanks. Sheet B) Enrichment test results for *copb2* regions with 1 kb flanks. Sheet C) Enrichment test results for common dragonet regions orthologous to the "seven gene loss" regions in syngnathids. Sheet D) Enrichment test results for the "seven gene loss" regions in the leafy seadragon genome.

**Supplemental File 5.** Annotations of miRNAs from leafy seadragon (peq) and weedy seadragon (pta) genomes, based on Illumina miRNA-seq data. The names, confidence in correspondence with other teleost miRNA annotations, locations, and sequences for the miRNAs are listed.
